# Speech markers of psychedelic-induced psychological change

**DOI:** 10.1101/2025.04.16.649217

**Authors:** Joanna Kuc, Rosalind G. McAlpine, Amelia Sellers, George Blackburne, Daniel R. Lametti, Jeremy I Skipper

## Abstract

5-MeO-DMT, a potent, short-acting psychedelic, induces profound shifts in cognition, affect, and self-awareness. Because language explicitly expresses these domains and voice implicitly conveys them, both may serve as potential ‘biomarkers’ of behavioural change. This study introduces a novel framework for analysing baseline language and vocal features, pre- to post-psychedelic changes, assessing their potential to predict subjective experiences and psychological outcomes. Daily voice journals from 29 participants were collected via ‘RetreatBot’ for two weeks before and after 5-MeO-DMT (1×12 mg). Transcripts were analysed using NLP (bag-of-words for vocabulary; transformer model for textual affect), and acoustic features (e.g., pitch, jitter, shimmer) were extracted to assess vocal dynamics. Following 5-MeO-DMT, speech markers revealed increased cognitive language, decreased social words, and altered voice quality (increased jitter/shimmer). Baseline speech patterns predicted psychological preparedness, ego dissolution anxiety, emotional breakthrough, and post-experience well-being. This first longitudinal analysis of speech markers following psychedelic use demonstrates a shift from external focus to introspection. Speech markers predicted and tracked psychological transformation, establishing vocal journaling as a valuable framework for monitoring psychedelic-induced changes and facilitating integration.

## Introduction

Psychedelics have reemerged in scientific and clinical research [1,2], including short acting compounds such as 5-Methoxy-N,N-dimethyltryptamine (5-MeO-DMT), which powerfully alter conscious experience and are currently evaluated for their potential to improve mental health [3,4]. Language is deeply tied to all human psychological domains, such as emotion and memory [5]. It has therefore been a useful tool for studying psychological functioning and mental health [6,7] in non-altered, everyday states of consciousness. Yet, there are currently no longitudinal studies examining ecologically produced language and voice characteristics for the purpose of understanding the mechanisms by which psychedelics have their effects. To fill this gap, we developed a conversational bot integrated into a mobile app to collect voice journals and used it to track participants before and after administration of 5-MeO-DMT.

### Understanding 5-MeO-DMT

The resurgence of interest in psychedelics has led to growing efforts to understand how these substances affect phenomenological experiences [8] and how this knowledge can be applied to mental health treatments [9]. The growing body of evidence suggests that, when administered in controlled settings, psychedelics can catalyse psychological transformation [8]. At the molecular level, psychedelics are associated with increased (neuro)plasticity [10], as well as changes in brain network dynamics [11,12]. At the experiential level, they alter fundamental aspects of consciousness (such as perception, self-awareness, and emotional processing [13]), with the intensity and quality of the subjective experiences predicting the long-term clinical outcomes [14].

5-MeO-DMT is a naturally occurring tryptamine, found in the secretions of the *Bufo alvarius (Incilius alvarius)*, the Sonoran Desert toad, and various plant species such as *Anadenanthera peregrina*, and also produced synthetically [15–17]. Compared to psilocybin or lysergic acid diethylamide (LSD), 5-MeO-DMT stands out for producing very intense but relatively brief alterations in consciousness, typically lasting 15–20 minutes when vaporised [3]. The subjective experience on 5-MeO-DMT is characterised by a rapid loss of sense of self and bodily awareness, often resulting in ‘mystical-type’ experiences, defined by a sense of oneness, intuitive or revelatory qualities, and an ineffable, paradoxical, and sacred nature, together with temporal and spatial distortions [17–21]. Distinctively, the 5-MeO-DMT experience is often described as “content-free”, featuring sensory deprivation frequently depicted as immersion in all-white light or total darkness [16]. Similarly to other psychedelics, 5-MeO-DMT is being evaluated for its potential beneficial long-term effects on mental health and well-being [3,4,16]. These outcomes are thought to be enhanced through adequate preparation prior to the session (psychoeducation, intention-setting, and establishing trust and comfort) integration afterwards (e.g. guided reflection and emotional processing) [22–24].

### Language as a tool

Modern views of complex psychological functions, like memory and emotional processing, suggest that they are supported by multiple networks, widely distributed throughout the whole brain [25]. This is also true of the neurobiology of language, perhaps even more so [5,26]. That is, humans use language ubiquitously, encountering about 150,000 words a day, beginning in utero [5]. These words are intimately tied to all of our psychological processes, including sensorimotor and social-emotional processing, and cognitive processes like attention and memory. For example, language is used to direct attention to and arrange the world categorically, organising sensory experiences into colours (e.g., ‘blue’), objects (‘banana’), and emotions (‘love’) [27–29]. It also organises our memories into ‘higher-level’ constructs like the ‘narrative self ‘ using words like ‘I’ and ‘me’ [30]. This explains why language has a broad neurobiological reach: Word processing is connected to widely distributed regions of the brain involved in all of these processes [5,26].

Because of its pervasive role in scaffolding a wide range of psychological processes, language may serve as a valuable proxy for understanding and predicting mental health [31]. Written diaries, for example, often capture preoccupations and facets of an individual’s inner dialogue, proving valuable in clinical psychology [32,33]. More recently, ecological momentary assessments (EMAs) - methodologies involving repeated sampling of experiences in real-world environments - have gained traction for their ability to capture naturalistic data from outside the typical research setting, and thus, minimise recall bias [34]. These EMAs can be administered through smartphone apps, prompting participants to respond to questionnaires or, increasingly, to provide open-ended text responses in real time. EMAs are now used both as a monitoring tool and as an intervention in mental health contexts [35], illustrating their versatility in capturing evolving linguistic markers of psychological well-being.

A common analytical approach to studying language use is the bag-of-words method, which treats language as a collection of individual words (without considering grammar or word order). Tools like Linguistic Inquiry and Word Count (LIWC), which captures the frequency of specific words in text, have demonstrated that word category frequencies can serve as indicators of various mental health conditions. For instance, depression correlates with increased use of absolutist terms and self-referential language [36–38], psychological wellbeing is reflected in patterns of emotional word use and cognitive processing terms [39], stress responses manifest in shifts in function words and emotional expression [40], and personality traits are marked by distinctive word-choice and grammatical structures [41]. While many of these findings stem from static text analyses, language-based EMAs expand these insights into real-world, real-time environments, e.g., aiding in the understanding of social aspects of alcohol consumption [42].

Alongside the word count approaches, the advent of transformer-based large language models (LLMs) has opened new frontiers in language analysis [43]. These models excel at contextual understanding and scaling to vast datasets, making them particularly powerful for capturing subtle semantic and pragmatic nuances. This is especially relevant in psychology, where factors such as register, metaphor, or sarcasm can reflect deeper emotional and cognitive states [6]. Recent applications of LLMs in mental health research include enhancing online psychological consultations [44], classifying mental health conditions like depression or anxiety [45], monitoring social media language for broader mental health surveillance [46], or detecting crisis signals in chat-based interactions [47]. Moreover, LLMs - which operate solely on language - can convincingly simulate diverse psychological profiles, leading users to empathise with them and perceive these models as psychologically similar to themselves [48]. These examples underscore the significant role of language in revealing psychological processes and the growing sophistication of language modeling methods in uncovering them.

In addition to lexical and semantic analyses, examining vocal features provides a complementary approach to understanding psychological states, offering objective, physiological markers that may reflect both acute and sustained changes in mental states [7]. Fundamental frequency, typically measured in Hertz (Hz), reflects emotional arousal and stress levels; jitter, measured as percent deviation from normal periodicity, indicates irregular changes in sound quality in a short time; and shimmer, reflecting irregular variation of amplitude [49]. These parameters have been linked to a range of psychological conditions: lower vocal variability, reduced pitch, and changes in loudness and tempo are associated with depression [50,51], while increased pitch variability has been observed in social anxiety disorder [52].

Nevertheless, methodologies across studies remain heterogeneous, both in analytical approaches and data sources (e.g., online posts, diaries, interviews). Given that language use evolves over time, across locations [53], and varies by contexts [54] and populations [55], the findings from one dataset may not generalise well to another [56], especially with small population sizes [57]. To ensure interpretability and relevance, language models must be attuned to the linguistic and contextual nuances of the populations they aim to represent.

### Language and psychedelics

Psychedelic experiences present a particularly compelling context for such tailored approaches, given their distinct effects on language and cognition. Building on the successful use of language-based methodologies to reveal underlying psychological processes, these techniques can likewise help us understand the mechanisms underlying these states and the mental health changes that may follow [58]. Supporting this, a large variety of studies have demonstrated that psychedelics alter language acutely, mostly by increasing ineffability [20,59,60] and changing word semantics to be less predictable [61–63], and more bizarre [64]. A number of studies have also used language to analyse reconstructed post-experience reflections (e.g., ‘trip reports’) and therapy transcripts to predict treatment outcomes.

Linguistic differences across substances have been identified through analysis of trip reports, with Hase et al. (2022) using a bag-of-words approach to uncover variations in analytical language and emotional expression within the Erowid corpus. Zamberlan et al. (2018) linked semantic similarities in trip reports to binding affinity profiles, bridging subjective language with pharmacology [65]. Žuljević et al. (2023) developed a mystical-experience dictionary, demonstrating correlations between mystical word usage and self-reported experience intensity [66]. Large-scale natural language processing of 11,000 Reddit trip reports revealed that psychedelics were more strongly associated with inferred emotions such as ‘Realization’, ‘Curiosity’, ‘Confusion’, ‘Surprise’, and ‘Amusement’ compared to other drug classes [67].

Beyond self-reports, sentiment analysis has been applied to therapy transcripts to predict treatment response. Carrillo et al. (2018) used a custom ‘emotional analysis’ algorithm with machine learning, achieving 85% accuracy in predicting psilocybin treatment response from 12 baseline interviews [68]. Dougherty et al. (2023) applied a transformer-based sentiment model to integration therapy sessions of 90 patients, achieving 85-88% accuracy in predicting long-term outcomes at three weeks post-dosing [69].

#### Current study

In summary, language is intimately connected to psychological functioning, making it a powerful marker for understanding psychological and mental health changes. There is clear evidence that language is impacted by psychedelics, though there seem to be few consistent results across studies. Existing work has focused on acute effects during the psychedelic state or analysed therapy sessions or post-experience reconstructions at a single time point, which may not capture language use in genuinely naturalistic settings and thus raise concerns about demand characteristics and ecological validity. Moreover, no prior studies have explicitly tracked longitudinal changes in natural language before and after psychedelic ingestion, and none have examined acoustic measures in these contexts - let alone tracked them from pre- to post-experience. Filling these gaps is crucial for uncovering objective markers of psychedelic-induced shifts in cognitive, affective, and social cognition, potentially shedding light on their mechanism of action and guiding more nuanced therapeutic applications. Thus, we conduct the first longitudinal investigation into both textual and acoustic features of natural speech surrounding a single high dose of 5-MeO-DMT. We focus on three key questions:

● **Pre- to post- 5-MeO-DMT language and voice changes:** How does vocabulary, emotions conveyed in language, and voice shift following 5-MeO-DMT exposure? What patterns emerge in emotional expression, and how do these changes relate to psychological outcomes?
● **Temporal Dynamics:** What patterns exist in language use across the study period? Are there systematic changes in vocal features before and after the experience, and can we identify markers that indicate successful preparation and integration?
● **Predictive Markers:** Can pre-experience language patterns predict experience intensity? Which linguistic features correlate with optimal outcomes, and how do vocal characteristics differ between preparation and integration periods?

Three main hypotheses guide this investigation. First, we predict that the psychedelic experience will induce detectable changes in language use, with certain linguistic shifts predicting psychological outcomes and subjective experience qualities. Second, we hypothesise that pre-experience language patterns, including both vocabulary usage and emotional expression, can predict the nature of the psychedelic experience and subsequent well-being outcomes. Third, we expect to observe distinct patterns in vocal features across different periods surrounding the psychedelic experience, reflecting phases of preparation and integration.

To test our hypotheses, we adopted an Ecological Momentary Assessment (EMA) approach using daily voice journals via a custom-built journaling bot (see Fig. 1). We analysed data collected within a ±14-day window around the single 12 mg 5-MeO-DMT session. We extracted bag-of-words features from transcribed speech using LIWC, classified text emotions with a RoBERTa model trained on the GoEmotions dataset, and performed acoustic analysis (e.g., pitch, jitter, shimmer) using the openSMILE library.

**Figure 1.**
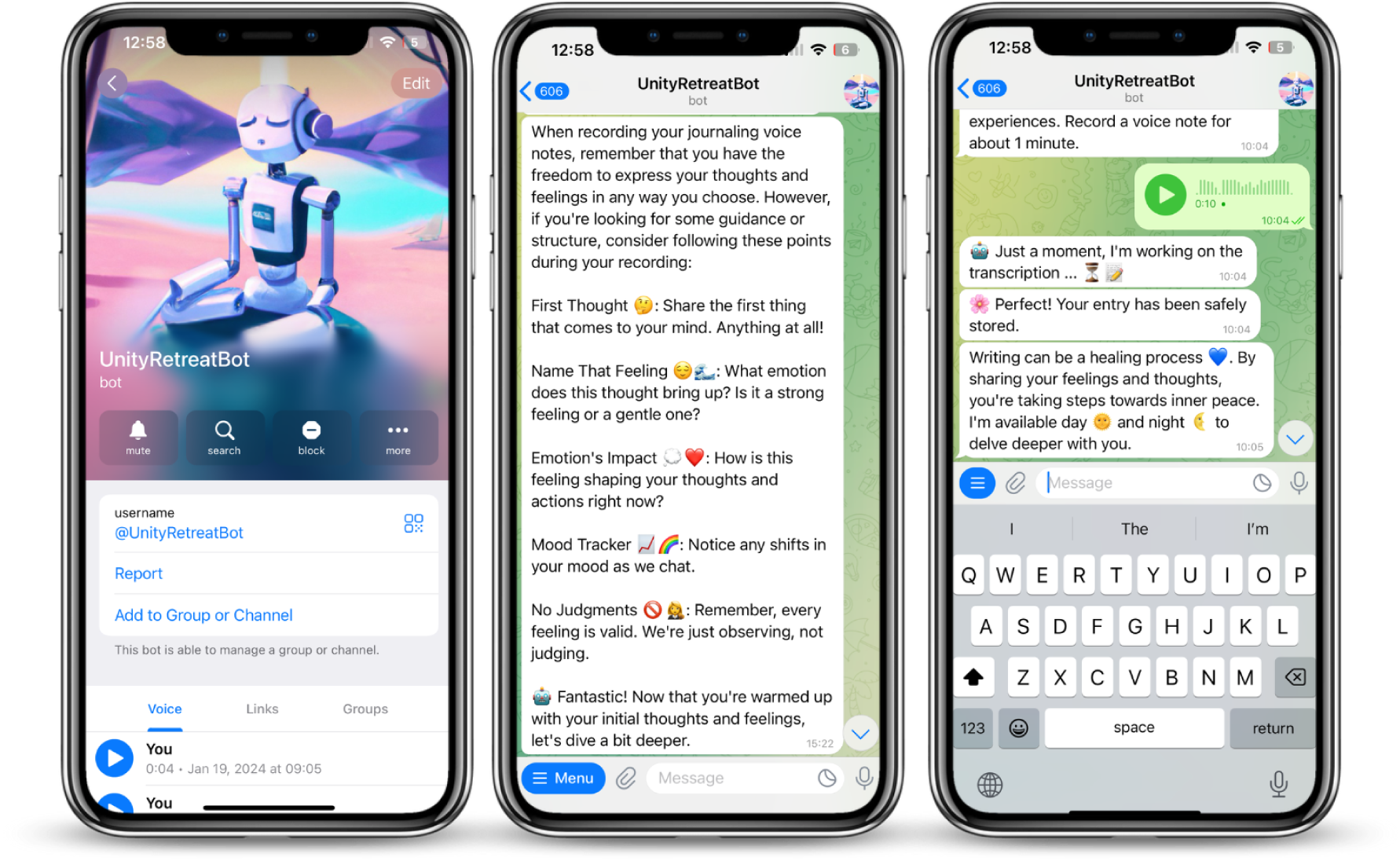
User interface of the ‘RetreatBot’ on Telegram. The left panel shows the bot’s profile screen where users initiate contact. The middle panel displays onboarding instructions introducing the journaling process and suggested reflection prompts. The right panel captures the journaling interaction flow, including a voice note recording, transcription, confirmation of entry submission, and a closing message of support.

## Results

### Daily submission patterns and participant engagement

Over a ±14-day period surrounding the 5-MeO-DMT session, participants (n=29) submitted a total of 288 unique voice journals comprising 4,127 sentences. These recordings represent 506.07 minutes (8.44 hours) of audio content. On average, each participant contributed 10 ± 6 voice journals and 142 ± 114 sentences (see: Fig. 2). The typical voice journal lasted 105.4 (±68.8) seconds. Submission frequency peaked shortly before the retreat, with heightened activity in the days immediately surrounding dosing. Although participants were prompted to submit entries from Days −7 to +7, we extended the analysis window to ±14 days based on observed engagement. Of the initial 29 participants, 25 provided both pre- and post-ceremony recordings, and 22 contributed at least two recordings in each period. Psychometric questionnaires were completed by 26 participants, with 24 providing at least two pre-dosing recordings plus questionnaire data. Sample sizes for each analysis are noted throughout. For more details on demographics, see Supplementary Table S1.

**Figure 2.**
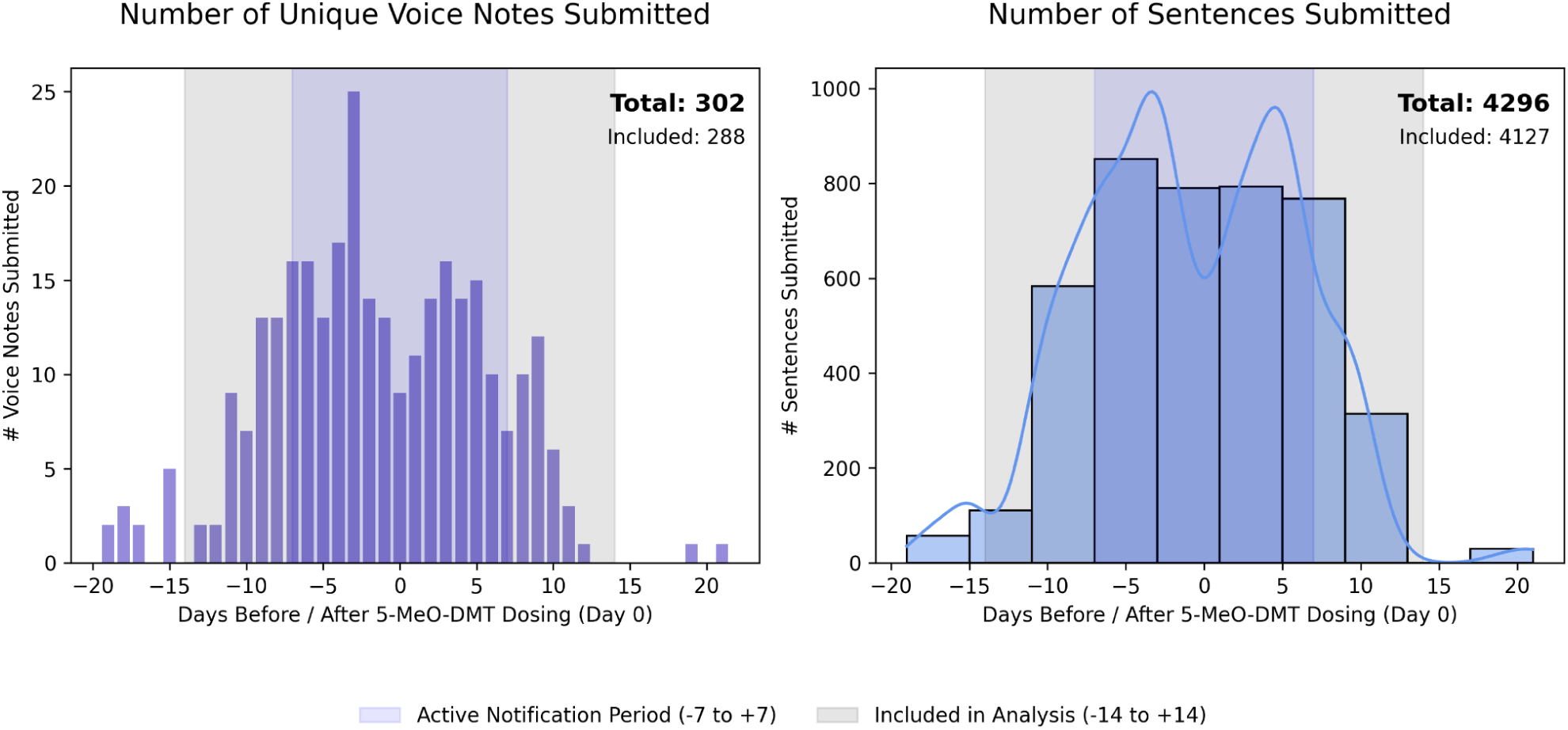
Daily submission patterns and participant engagement before and after 5-MeO-DMT dosing. The bars on the left panel represent the total number of unique voice journals submitted per day relative to the 5-MeO-DMT dosing session (Day 0). The right panel displays the distribution of sentences submitted per day, with blue bars representing the total number of sentences and the blue line showing the Kernel Density Estimation (KDE) to visualise the smoothed distribution pattern. The light blue shaded region (−7 to +7 days) represents the active notification period, during which participants were actively reminded to submit entries. The light grey shaded region (−14 to +14 days) represents the full period included in the analysis. Total values for both unique entries and sentences are indicated in the top right corner of each panel.

### Textual shifts following 5-MeO-DMT

To examine individual changes (n=29) in vocabulary use from before to after the 5-MeO-DMT experience, we analysed the LIWC categories that reflect linguistic, psychological, social aspects of language.

Overall, a number of significant linguistic shifts across multiple LIWC categories were observed in participants’ voice journals after the 5-MeO-DMT experience (Fig. 3, left panel; for full statistical results see: Supplementary Table S2). *Cognitive* word use increased significantly, (Coef.: 1.96, p_FDR_ < 0.001), driven by greater frequency of *cognitive processes* words (e.g., “because”, “felt”, “pretty”, “if”, “want”; Coef.: 1.93, p_FDR_ < 0.001), *insight* words (e.g. “felt”, “know”, “feeling”, “reflecting”; Coef.: 0.63, p_FDR_ < 0.05), *tentative* words (e.g., “pretty”, “if”, “any”, “sometimes”, “hope”, “might”; Coef.: 0.74, p_FDR_ < 0.01), and *certitude* words (e.g., “really”, “real”, “actually”, “incredibly”, “totally”; Coef.: 0.43, p_FDR_ < 0.05).

**Figure 3.**
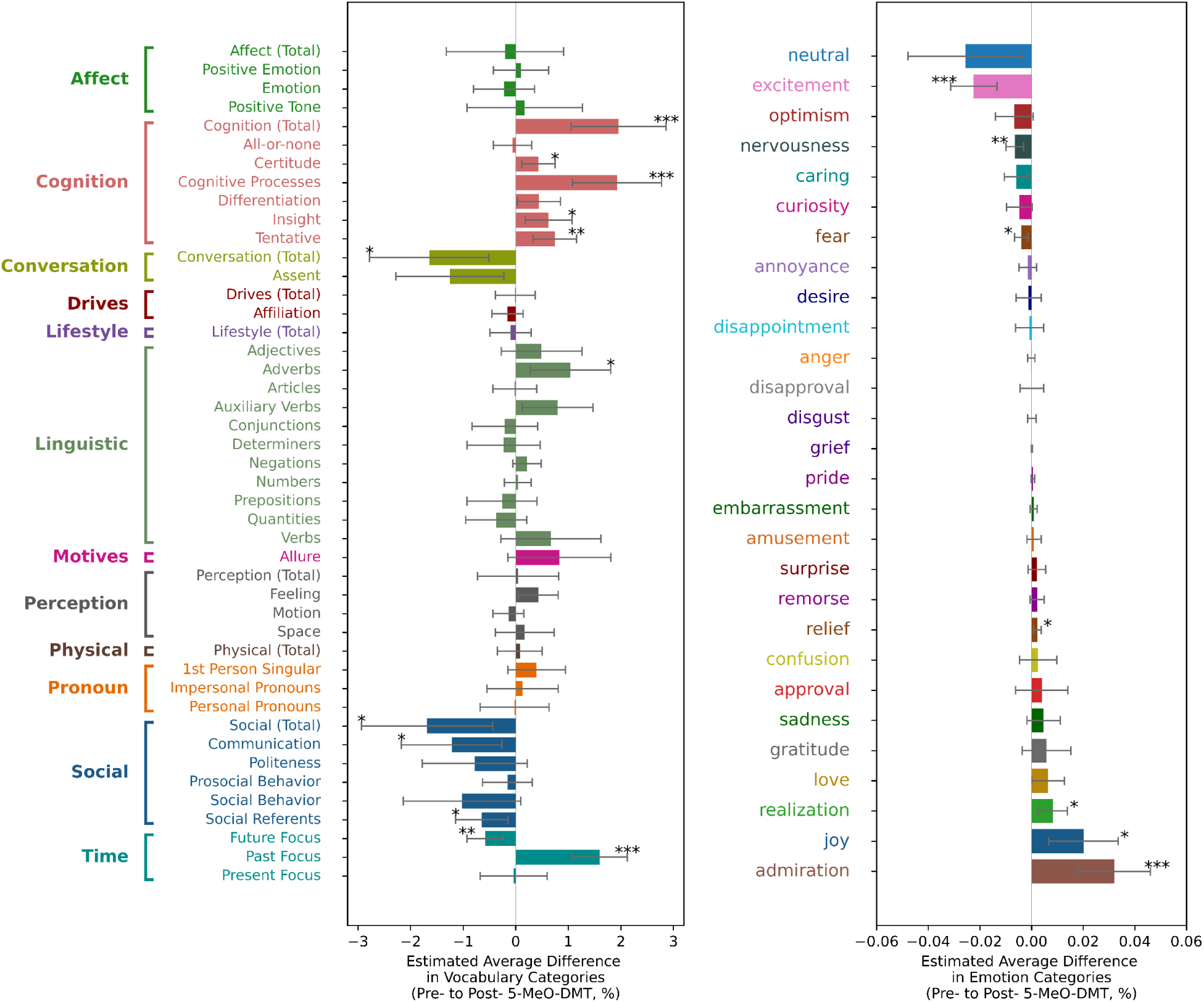
Changes in textual metrics from pre- to post-5-MeO-DMT dosing. Mixed-effects model estimates of sentence-level changes (n=4127 sentences, 29 participants) in LIWC vocabulary categories (left) and RoBERTa-GoEmotions probabilities (right). Each bar represents the intercept from a separate mixed-effects model with a binary “Pre-Post” predictor variable (0/1), with 95% confidence intervals (feature ∼ PrePost + 1/ParticipantID). All p-values FDR-corrected (Benjamini-Hochberg). p < 0.05, p < 0.01, p < 0.001.

By contrast, *Social* words (e.g., “they”, “life”, “she”, “we’re”, “he’s”; Coef.: −1.69, p_FDR_ < 0.05) decreased, with most reductions seen in the *communication* (e.g., “thank”, “talk”, “meeting”, “say”, “said”, “told”; Coef.: −1.22, p_FDR_ < 0.05) and *social referents* (e.g., “they”, “she”, “friends”, “everybody”; Coef.: −0.64, p_FDR_ < 0.05) subcategories.

In the *Linguistic* category, the use of *adverbs* increased significantly (e.g., “pretty”, “when”, “so”, “now”, “there”, “here”; Coef.: 1.04, p_FDR_ < 0.05), while *conversational markers* declined (e.g., “um”, “okay”, “yes”, “yeah”.; Coef.: −1.64, p_FDR_ < 0.05). Within the *Time* category, there was a significant increase in *past-oriented language* (e.g., “went”, “made”, “yesterday”, “realised”; Coef.: 1.6, p_FDR_ < 0.001), and a significant decrease in *future-oriented language* (e.g., “tomorrow”, “ready to”, “hoping”, “we’ll”, Coef.: −0.58, p_FDR_ < 0.01).

Text-based emotion detection using a RoBERTa-GoEmotion model (see Fig. 3, right panel; for full statistical results see: Supplementary Table S3) revealed that after 5-MeO-DMT administration, participants (n=29) showed significantly decreased *excitement* (Coef.: −0.022, p_FDR_ < 0.001), *nervousness* (Coef.: −0.007, p_FDR_ < 0.01), and *fear* (Coef.: −0.004, p_FDR_ < 0.05), while *admiration* (Coef.: 0.028, p_FDR_ < 0.01), *relief* (Coef.: 0.003, p_FDR_ < 0.01), *joy* (Coef.: 0.022, p_FDR_ < 0.05), and *realisation* (Coef.: 0.009, p_FDR_ < 0.05) increased.

#### Vocal feature changes

Three of 88 vocal features derived from openSMILE (n=23) demonstrated significant post-dosing alterations in jitter and shimmer. Average local jitter (Coef. = 0.776, p_FDR_ < 0.05) and shimmer (Coef. = 0.712, p_FDR_ < 0.05) increased, while normalised jitter variability decreased (Coef. = −0.910, p_FDR_ < 0.05). Full statistical results are provided in Supplementary Table S4 and visualised in Supplementary Figure S3.

#### Temporal dynamics of cognitive and social language

Consistent with overall linguistic analysis (see Fig. 3), the day-by-day analyses (n=25; see Fig. 4) showed a pronounced crossover in cognitive versus social word usage after the 5-MeO-DMT session. Whereas *social* language, initially above baseline, fell below baseline levels post-dosing, *cognitive* language rose above baseline. This switch in trajectories persisted into the second post-dosing week, reflecting a sustained transition from socially oriented discourse toward a more cognitively focused language profile following the psychedelic experience (see Fig. 4, top). Word clouds revealed characteristic terms in cognitive language (e.g., “because”, “felt”, “if”, “want”, “trying”, “know”) and social language (e.g., “they”, “life”, “she”, “we’re”, “talk”, “city”, “friends”) that drove these opposing trajectories (see Fig. 4, middle). This pattern was confirmed statistically, with a significant increase from weeks −1 to 1 and 2 for cognitive words and a significant decrease in social words from weeks −1 to 1 (see Fig. 4, bottom). For detailed statistics see Supplementary Tables S5 and S6.

**Figure 4.**
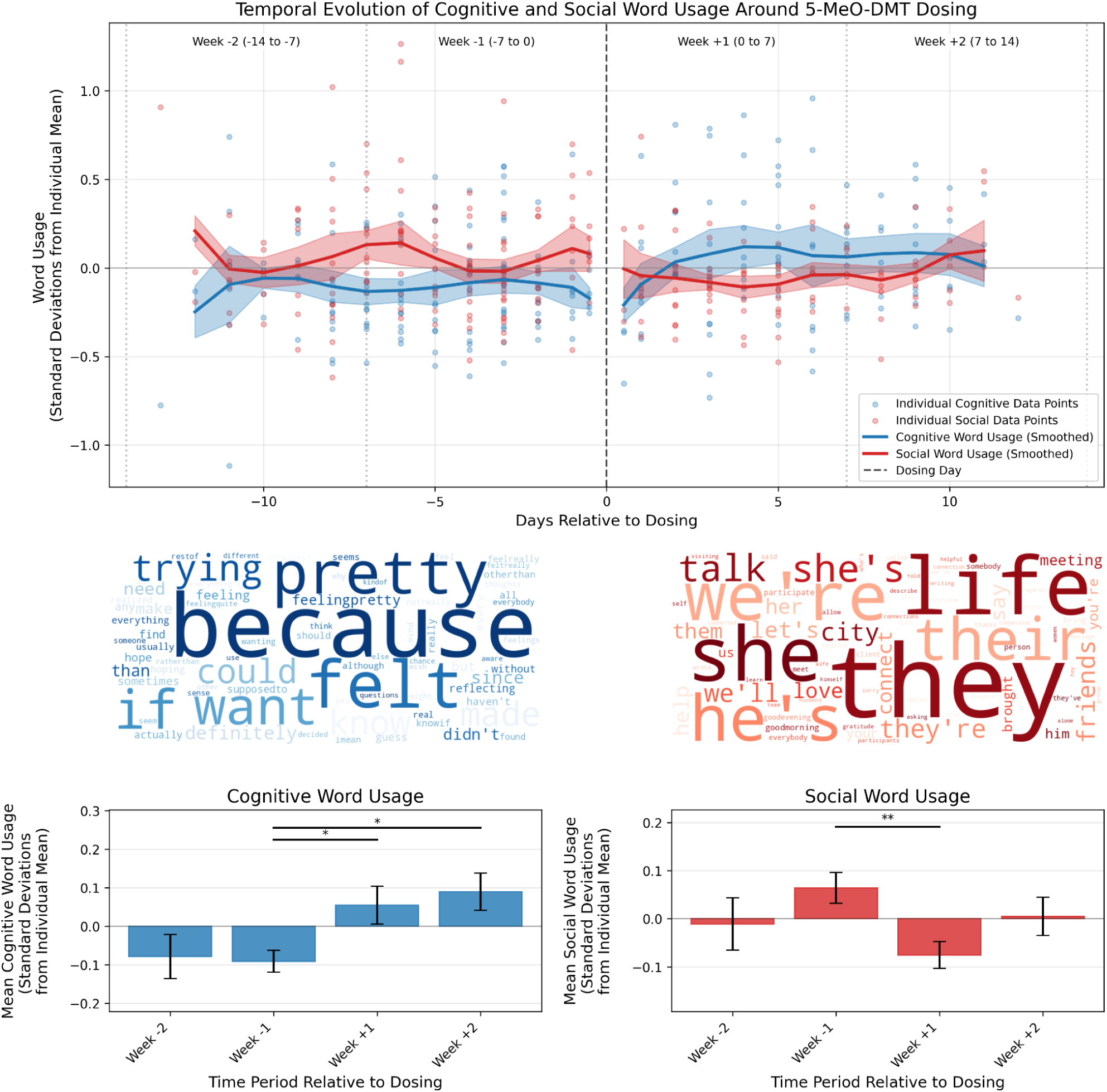
Temporal evolution of cognitive and social word usage around 5-MeO-DMT intervention. Top panel: Time series showing standardised deviations from individual means in cognitive (blue) and social (red) word usage from 14 days before to 14 days after dosing (n=25 participants; 4133 sentences analysed). Light-colored points represent individual data points, while solid lines show Gaussian-smoothed averages (σ=1) averages with standard error bands. Background colors indicate weekly periods used for statistical comparisons. Middle panels: Word clouds showing the most frequently used cognitive (left, purple) and social (right, green) vocabulary terms. Bottom panels: Weekly averages of (left) cognitive and (right) social word usage with 95% confidence intervals. Statistical significance bars indicate differences between periods based on independent t-tests with after FDR correction; * p < 0.05, ** p < 0.01, *** p < 0.001.

#### Temporal dynamics of vocal feature deviations

Pitch, jitter, and shimmer - selected for their associations with mental health and emotional regulation - showed distinct shifts around the 5-MeO-DMT session (n=22; see Fig. 5). For each feature, deviations from individual baselines (±1 SD) were classified as “high” or “low” and tracked daily. Day-by-day analyses (Fig. 7, top) revealed that *high pitch* deviations peaked at Day −8 (frequency = 5.28), whereas *low pitch* deviations peaked at Day +2 (frequency = 2.98). Group-level comparisons across predefined weekly periods (Fig. 7, bottom) confirmed a significant reduction in high pitch deviations from Week −1 (frequency = 2.23) to Week +2 (frequency = 0.51; pFDR = 0.0353), and a significant increase in low pitch deviations from Week −2 (frequency = 0.76) to Week +1 (frequency = 1.98; pFDR = 0.0079). Similar patterns were observed for jitter and shimmer, with elevated low deviation frequencies during the post-session period.

**Figure 5.**
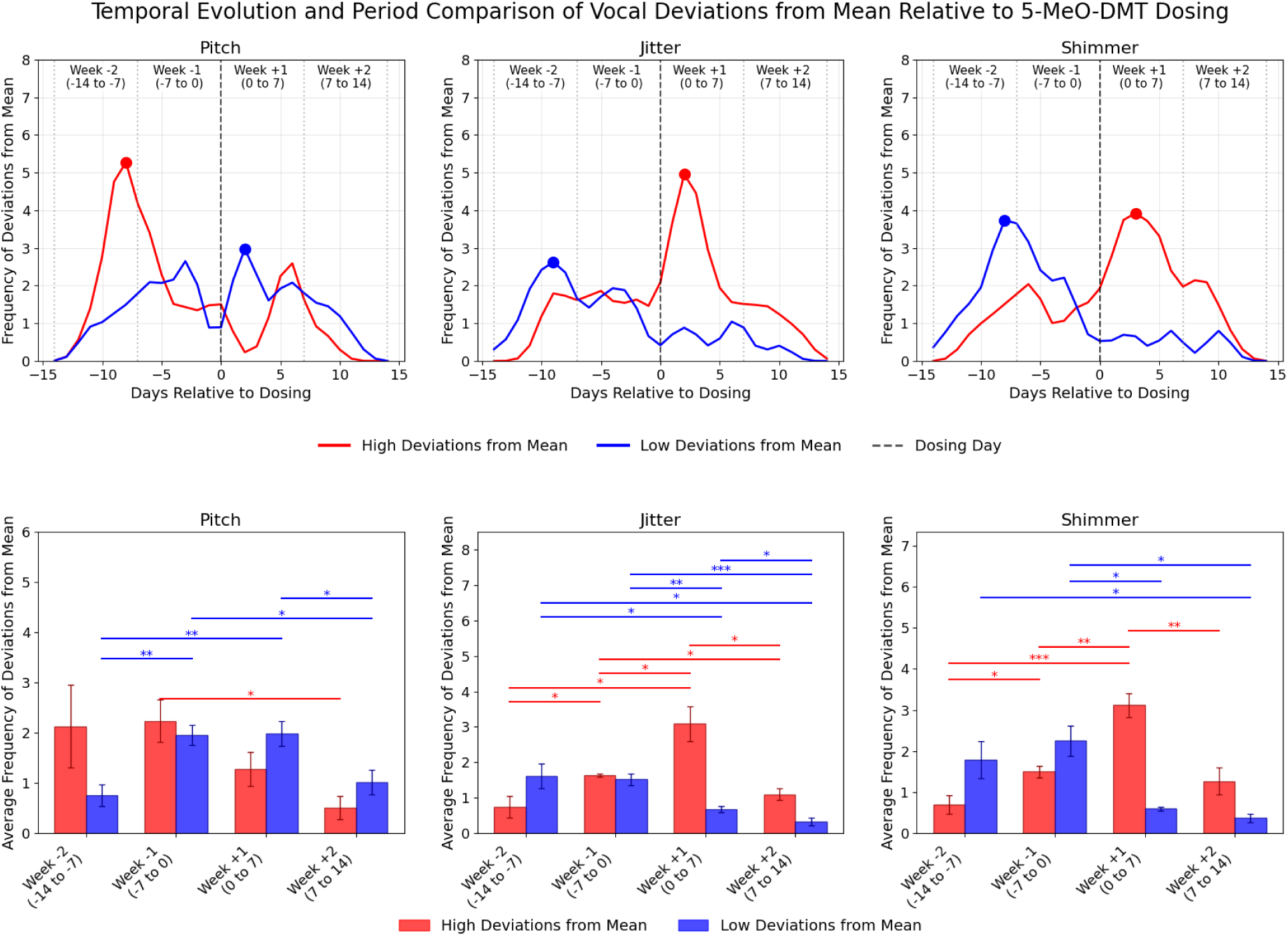
Temporal Evolution and Period Comparison of Vocal Deviations from Mean Relative to 5-MeO-DMT Dosing. (Top) Line plots showing the smoothed daily frequency of high (red) and low (blue) deviations in pitch, jitter, and shimmer over a 28-day period, centered around the dosing day (Day 0, vertical dashed line). Deviations were classified as significant when participants’ vocal measurements exceeded one standard deviation above (high) or below (low) their individual mean values. Vertical dotted lines demarcate weekly periods: Week −2 (−14 to −7 days), Week −1 (−7 to 0 days), Week +1 (0 to 7 days), and Week +2 (7 to 14 days). Colored dots indicate peak frequency points for high and low deviations, with smoothing applied using a Gaussian filter (sigma=1). (Bottom) Bar plots depicting mean frequency of smoothed deviations during each weekly period, with error bars representing standard error of the mean (SEM). Statistical significance between periods is based on independent t-tests with False Discovery Rate (FDR) correction for multiple comparisons (*p < 0.05, **p < 0.01, ***p < 0.001). Only participants with at least two recordings in both pre and post periods were included in the analysis.

Jitter presented the opposite pattern, with *high jitter* deviations peaking at Day +2 (frequency = 4.95) and *low jitter* deviations at Day −9 (frequency = 2.63). Over weekly intervals, high jitter deviations rose from Week −2 (frequency = 0.74) to Week −1 (frequency = 1.63; pFDR = 0.0320) and subsequently to Week +1 (frequency = 3.09; p_FDR_= 0.0147). Low jitter deviations decreased from Week −2 (frequency = 1.61) to Week +1 (frequency = 0.68; p_FDR_= 0.0486) and further to Week +2 (frequency = 0.33; pFDR = 0.0213).

Finally, *high shimmer* deviations peaked at Day +3 (frequency = 3.92), whereas *low shimmer* deviations were most frequent at Day −8 (frequency = 3.73). Weekly comparisons showed shimmer high deviations rising from Week −2 (frequency = 0.69) to Week −1 (frequency = 1.50; p_FDR_= 0.0175) and to Week +1 (frequency = 3.12; p_FDR_= 0.0002), before falling back by Week +2 (frequency = 1.27; p_FDR_= 0.0025). Shimmer low deviations decreased from Week −1 (frequency = 2.25) to Week +1 (frequency = 0.60; p_FDR_= 0.0124) and further to Week +2 (frequency = 0.37; p_FDR_= 0.0112).

We conducted an additional analysis, to examine the temporal dynamics of vocal features in a less categorical manner, at the individual-level and using the same methods as those applied for the vocabulary dynamics (see: Supplementary Fig. S2). This confirmed the main temporal trends in pitch and jitter observed at the group level. Shimmer followed a similar trajectory but did not reach corrected statistical significance.

### Prediction of psychometric outcomes from language features

#### Pre-dosing vocabulary patterns as predictors of questionnaire responses

We examined correlations between weighted means of pre-dosing LIWC features and five psychometric measures using a moderately strong threshold of |r| > 0.6, which served as our initial feature selection approach, in order to retain only the strongest linguistic predictors for subsequent regression analyses. All analyses were conducted with 26 participants after removing cases with missing data. We focused on five key measures to capture core aspects of the psychedelic experience and its impact on well-being: preparedness (Psychedelic Preparedness Scale, PPS), negative ego dissolution (Altered States of Consciousness – Dread of Ego Dissolution subscale, ASC-DED), unitive experience (Altered States of Consciousness – Oceanic Boundlessness subscale, ASC-OBN), emotional breakthroughs (Emotional Breakthrough Inventory, EBI), and mental well-being (short Warwick–Edinburgh Mental Well-Being Scale, sWEMWBS). Together, these domains reflect both immediate experiential qualities and post-experience psychological outcomes.

Strong correlations were found between the PPS and three linguistic features: positive tone vocabulary (e.g.: “good”, “well”, “beautiful”, “thank”, “excited”; r = 0.654), work-related language (e.g., “work”, “session”, “meeting”, “study”, “school”; r = 0.636), and lifestyle references (e.g., “work”, “home”, “spent”, “bed”, “session”, “weekend”; r = 0.612). ASC-DED subscale correlated with quantity-related language (e.g., “day”, “whole”, “week”, “most”, “few”; r = 0.628), while sWEMWBS negatively correlated with mental health-related language (e.g., “psychologist”, “delusion”, “mental health”, “suicide”; r = −0.617). No strong correlations were identified for the EBI or the ASC-OBN scale.

Linear regression models were constructed separately for each psychometric measure (dependent variable) using its strongly correlated LIWC features as predictors (independent variables). For the PPS outcome model (R² = 0.575, F(3,22) = 9.920, p < 0.001), only positive tone remained a significant predictor (β = 2.492, p_FDR_ = 0.014), while work-related language (β = 5.343, p_FDR_ = 0.333) and lifestyle references (β = 0.562, p_FDR_ = 0.850) did not reach significance. The ASC-DED outcome model (R² = 0.394, F(1,24) = 15.634, p < 0.001) confirmed quantity-related language as a significant predictor (β = 0.105, p_FDR_ = 0.002). Similarly, the sWEMWBS outcome model (R² = 0.380, F(1,24) = 14.727, p < 0.001) validated mental health-related language as a significant negative predictor (β = −34.110, p_FDR_ = 0.002).

To evaluate model robustness and capture broader multivariate contributions, we additionally conducted ridge regression analyses using all weighted means of pre-dosing LIWC features as predictors. Results largely aligned with findings from the linear models. The PPS model (R² = 0.580) identified work-related language (β = 0.331), lifestyle references (β = 0.315), and positive tone (β = 0.284) as the strongest predictors, consistent with correlation-based selection. Similarly, for sWEMWBS (R² = 0.417), mental health-related language (β = −0.098) emerged as the strongest negative predictor, with additional features—such as lifestyle references (β = 0.077), cognition (β = −0.069), cognitive processes (β = −0.066), and leisure-related language (β = 0.062)—contributing minimally. The ASC-DED model showed high predictive performance (R² = 0.872), but the top features—including quantity-related language (β = 0.020), sexual references (β = 0.027), and future focus (β = −0.020)—had negligible coefficients, suggesting low interpretability despite strong model fit. For outcomes not associated with strong effects in the linear models, ridge regression identified additional predictive features. The EBI model (R² = 0.730) was driven by risk-related language (β = 2.177), illness references (β = 2.125), netspeak (β = −1.906), and curiosity-related terms (β = −1.781), all showing large coefficients. In contrast, the ASC-OBN model (R² = 0.508) yielded uniformly small coefficients for top predictors, such as substance references (β = −0.011) and friend-related language (β = −0.010), limiting the interpretability of that model. Full results are reported in Supplementary Table S7.

#### Pre-dosing emotional states and their relationship to outcome measures

Principal Component Analysis (PCA) performed on the pre-dosing weighted mean emotion scores derived from the RoBERTa-GoEmotions model identified three key patterns of emotional expression combinations, together explaining 49.1% of the total variance (see: Supplementary Fig. S3). Each of the identified Principal Components (PCs) was then used as a predictor in multiple regression analyses to examine their relationship with selected psychological outcomes (for details see: Supplementary Table S8).

**PC1** explained 25.5% of the variance and we refer to it as ‘positive emotional engagement’ (see: Fig. 6, top). This is because it represented a contrast between *positive emotional expression*, characterised by gratitude (0.303), pride (0.266), joy (0.258), excitement (0.234), caring (0.223), and love (0.206), versus *negative emotional states*, marked by disapproval (−0.282), confusion (−0.264), embarrassment (−0.257), disgust (−0.250), disappointment (−0.245), and annoyance (−0.235). When mapping vocabulary categories onto PC1, we found significant positive correlations with positive tone words (r = 0.596, p_FDR_ = 0.032) and significant negative correlations with negations (e.g., “didn’t”, “haven’t”, “not really”, “not”; r = −0.658, p_FDR_ = 0.011), cognition (r = −0.743, p_FDR_ = 0.001), cognitive processes (r = −0.748, p_FDR_ = 0.001), and differentiation (e.g., “if”, “didn’t”, “than”, “but”, “else”, “without”; r = −0.647, p_FDR_ = 0.012). As shown in the bottom panel of Fig. 6, linear regressions confirmed that higher PC1 scores significantly predicted greater preparedness as measured by the PPS (β = 2.33, p = 0.002, p_FDR_ = 0.017, R² = 0.363) and sWEMWBS (β = 0.66, p = 0.005, p_FDR_ = 0.020, R² = 0.303) scores.

**Figure 6.**
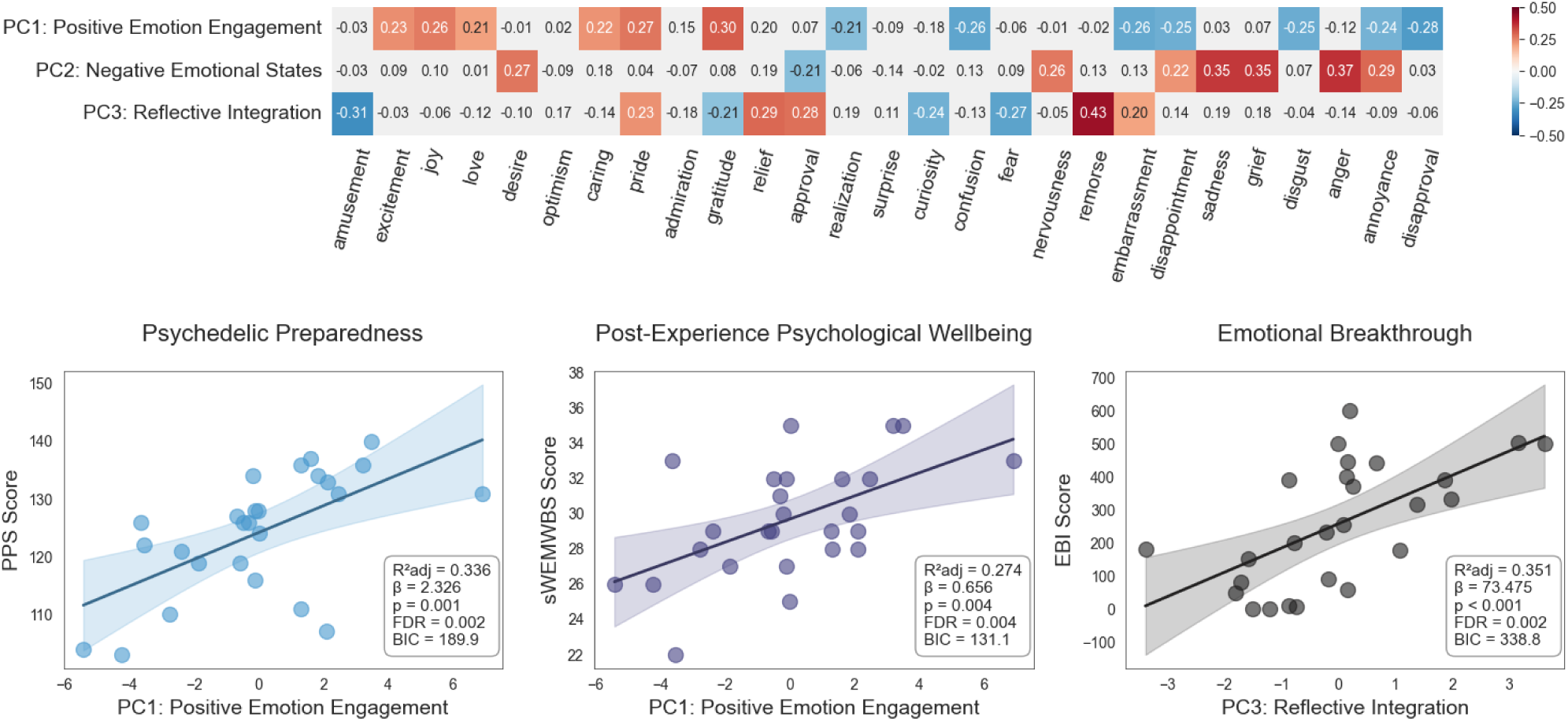
Principal component analysis of emotions and their relationships with psychological measures. Top panel: Heatmap of principal component loadings for the first three components. Emotions were detected using the RoBERTa-GoEmotions model to rate individual sentences in participants’ pre-dosing narratives. Weighted mean emotion scores were calculated for each participant and subjected to principal component analysis. Colors represent loading strengths (red = positive, blue = negative), with intensity indicating magnitude. Values show correlation Coef.s between emotions and components (PC1-PC3). Bottom panel: Linear regression analyses examining the relationships between the principal component and three psychological measures: psychedelic preparedness, post-experience psychological wellbeing, and emotional breakthrough. Shaded areas represent 95% confidence intervals. Statistics shown include standardized beta Coef.s (β), p-values (p), and R-squared values (R²).

**PC2** (15.3% variance), referred to as ‘negative emotional states’ contrasted intense emotional distress, including anger (0.366), sadness (0.355), grief (0.353), annoyance (0.288), desire (0.267), nervousness (0.263), and disappointment (0.224), with emotional neutrality (−0.317) and approval (−0.212). When mapping LIWC categories onto PC2, we found significant positive correlations with 3rd person singular (r = 0.639, p_FDR_ = 0.019), negative tone (e.g., “tired”, “nervous”, “lost”, “anxiety”, “sick”; r = 0.659, p_FDR_ = 0.016), emotion (e.g., “good”, “tired”, “excited”, “happy”, “nervous”; r = 0.613, p_FDR_ = 0.027), negative emotion (e.g., “tired”, “nervous”, “anxiety”, “sad”; r = 0.601, p_FDR_ = 0.029), and anger (e.g., “frustrating”, “angry”, “revenge”, “pissed off”; r = 0.725, p_FDR_ = 0.004). PC2 did not predict any of the outcome variables.

**PC3** (8.3% variance), ‘reflective integration’, captured the tension between reflective emotions, expressed through remorse (0.427), relief (0.287), approval (0.276), pride (0.230), and embarrassment (0.203), versus exploratory emotions characterised by amusement (−0.309), fear (−0.267), curiosity (−0.242), and gratitude (−0.206). No significant correlations were found when mapping LIWC categories onto PC3 after FDR correction. However, higher PC3 scores predicted a stronger emotional breakthrough, as measured by the EBI (see: Fig. 6. bottom panel; β = 73.47, p = 0.001, p_FDR_ = 0.010, R² = 0.377).

## Discussion

Our investigation into participants’ naturalistic language and voice surrounding a single 5-MeO-DMT session provides the first comprehensive demonstration that psychedelic experiences predict measurable longitudinal changes in verbal and vocal expression, revealing psychologically meaningful dynamics in real-world contexts. By capturing voice journals for two weeks before and after dosing, we identified significant shifts in cognitive and social language use, observed altered linguistic markers of emotional expression, and uncovered distinct vocal markers - together pointing to a reorientation of participants’ internal and external focus. These findings suggest that linguistic and vocal markers may reflect core psychological mechanisms of integration and offer potential predictors of therapeutic response. This study also introduces a set of methodological innovations: the use of voice-note diaries as an ecological momentary assessment tool, a custom chatbot for automated longitudinal data capture, and a multimodal analytical framework combining bag-of-words vocabulary analysis, transformer-based textual emotion detection, and acoustic feature extraction. These approaches enabled fine-grained, low-burden monitoring of psychological states in the periods leading up to and following the psychedelic experience (often termed ‘preparation’ and ‘integration’ in therapeutic contexts), and offer a scalable framework for future clinical and naturalistic research.

### Linguistic and vocal indicators of post-session integration

A key finding was the marked increase in cognitive and decrease in social words following 5-MeO-DMT, suggesting a reorientation from externally focused discourse toward more introspective modes of expression (Fig. 4, left). Day-by-day trajectories (Fig. 6) showed a crossover effect: social language, initially elevated relative to baseline, declined sharply post-dosing, while cognitive language rose and remained elevated for up to two weeks. Such a pattern points to a cognitively active, “slightly detached” phase that may persist during the two weeks after dosing, as participants work to contextualise and process the psychedelic experience. Despite the absence of formal integration therapy in this study, this interval may still support heightened self-examination and meaning-making. This process may relate to post-dosing changes in brain function, including increased neuroplasticity in regions like the prefrontal cortex, which is involved in emotional regulation [10,70]. Indeed, increased use of cognitive process terms (e.g., *because*, *felt*, *pretty*, *if*, *want*) and cognitive differentiation vocabulary (e.g., *if*, *didn’t*, *than*, *but*, *else*, *without*) may reflect intensified reflective thinking or problem-solving. This aligns with evidence that psychedelics can enhance cognitive flexibility [71] and suggests that language could serve as a useful marker for tracking such processes.

While the overall number of affective words and the balance between positive and negative emotion terms (as measured by a bag-of-words approach; Fig. 4, left) remained relatively stable from pre- to post-dosing, a more nuanced analysis using a transformer-based model (Fig. 4, right) reveals a substantial shift in the *specific types* of emotions verbally expressed across timepoints. Emotions related to anticipation and uncertainty (e.g., excitement, nervousness, fear) gave way to expressions reflecting positive affect and meaning-making (e.g., joy, admiration, realisation), suggesting a shift from *looking forward* to *looking inward*. Notably, the rise in “realisation” aligns with theories that psychedelics catalyse insight and perspective shifts [72,73], pointing toward an active process of integrating and making sense of the experience rather than a simple elevation of mood. These text-inferred affective changes, coincided with a significant increase in past-focused language and a decrease in future-oriented expression, suggest a shift from anticipatory thought to retrospective processing. Together with reduced social and increased cognitive word use, this pattern points to a post-acute phase of introspective engagement. This pattern may reflect a temporary distancing from interpersonal concerns and anticipatory thinking, replaced by a focus on immediate, introspective, and emotionally salient content.

Alongside these linguistic dynamics, we observed significant shifts in vocal parameters-particularly jitter and shimmer-that offer further insight into the aftermath of 5-MeO-DMT. Both jitter and shimmer increased on average from pre- to post-session, while jitter variability decreased, reflecting a distinctive pattern of amplitude and frequency modulation (Fig. 5). Although higher jitter and shimmer have been linked to anxiety [74–76] and depression [77], the absence of elevated anxiety (see: Supplementary Table S1) in our dataset suggests that these changes may reflect a temporary “turbulent” or liminal state, rather than pathology. This interpretation is exploratory and draws tentative support from emerging, primarily preclinical, evidence that psychedelics can reopen ‘critical periods’ of neuroplasticity [78], which could allow for the revision of entrenched cognitive and emotional patterns. In our day-by-day analyses over a 28-day window, pitch was frequently elevated prior to dosing - a “build-up” phase - while jitter and shimmer reached their highest deviations in the immediate post-session period, implying a subsequent “integration” phase of sustained emotional engagement (Fig. 7). Although additional single-participant analyses confirmed the main pitch and jitter trends, shimmer showed more variability across individuals, suggesting personal differences in amplitude modulation. These vocal changes, when considered alongside shifts in cognitive and affective language use, highlight a multifaceted process of ‘transition’ following 5-MeO-DMT. While participants’ textual narratives increasingly focused on introspection and meaning-making, elevated jitter and shimmer may reflect an embodied, physiological dimension of heightened engagement with emotional and cognitive content.

Taken together, these findings underscore that elevated jitter and shimmer might not necessarily indicate clinical distress but rather a transient window of malleability and openness following a profound psychedelic event. Importantly, this divergence between surface-level emotional stability and deeper shifts in emotional tone highlights the complementary strengths of different analytic approaches. While bag-of-words methods capture overall linguistic trends, transformer-based models uncover subtle context-dependent transformations in emotional expression, and the observed changes in vocal parameters reveal additional psychophysiological nuances. Taken together, these convergent measures shed new light on the extended psychological processes that follow a single 5-MeO-DMT session and underscore the value of multimodal approaches in understanding psychedelic experiences.

#### Language-based insights into self-reported outcomes

Our findings also reveal a nuanced relationship between participants’ pre-dosing language use and the outcomes of their psychedelic sessions, beginning with psychedelic preparedness. Participants who verbally expressed more positive emotions pre-dosing - captured by PC1-also used more positive tone language (e.g., *good*, *thank*, *excited*) and fewer negations, cognitive terms, and differentiation words. Both PC1 and positive terms independently predicted higher PPS scores, suggesting that emotionally affirmative language use reflects greater preparedness for experience. A similar pattern emerged for self-reported well-being (sWEMWBS), which was positively associated with PC1 and negatively associated with mental health-related vocabulary (e.g., *psychologist*, *delusion*, *mental*, *suicide*). This suggests that language with a positive tone may serve as both an indicator and facilitator of psychological readiness, thereby enhancing therapeutic outcomes in psychedelic interventions.

An intriguing finding arose in predicting emotional breakthrough, a state of *emotional catharsis* often linked with beneficial therapeutic outcomes in psychedelic-assisted treatments for depression [69,79,80]. While no specific vocabulary clusters were associated with EBI scores, higher PC3 scores-reflecting greater expression of reflective emotions such as remorse, relief, approval and embarrassment-significantly predicted stronger emotional breakthroughs. Notably, these emotions are not uniformly positive; rather, they suggest a capacity to engage with complex or ambivalent inner states. This supports the view that openness to emotionally challenging or integrative content may facilitate deeper therapeutic processing [79]. It also highlights how some experiential outcomes may elude capture by discrete vocabulary categories, underscoring the value of transformer-based approaches in detecting more subtle emotional dynamics. Finally, language use also appeared to reflect the negative expectation or apprehension toward ego dissolution. Greater use of quantity-related expressions (e.g., *day*, *whole*, *week, little bit, most*) in pre-dosing language was associated with heightened self-reported post-experience dread of ego dissolution. This pattern may reflect a more structured, time-bound cognitive style, with the focus on duration or magnitude signalling anxiety about losing control or self-coherence.

In summary, these findings support emerging models of psychedelic preparedness, highlighting how baseline emotional and linguistic patterns shape participants’ readiness and their subsequent experiences. Rather than relying solely on retrospective self-report, natural language offers a dynamic, unobtrusive window into the psychological state-one that may help identify individuals who could benefit from tailored preparation. Ongoing, voice-based assessments may further illuminate how readiness and risk evolve over time, ultimately enabling more responsive support before, during, and after psychedelic sessions.

### Limitations

A key limitation of this study is its sample composition: a small, self-selecting group of individuals with extensive prior experience using psychedelics, who are not representative of the general population [81,82]. Psychedelic users tend to exhibit higher openness and extraversion, and lower neuroticism [83,84], traits that may influence spoken language complexity and expressivity [85]. Moreover, participants’ high familiarity with 5-MeO-DMT (mean lifetime use: 39 occasions; SD = 72.91, range = 1–300) may have also influenced baseline linguistic patterns, potentially obscuring session-specific effects. Because all participants were healthy volunteers without diagnosed mental health conditions, the generalisability of these findings to clinical populations remains limited-an important consideration given the increasing use of psychedelics in therapeutic contexts [86], where language [87] and speech [88] are being studied as potential diagnostic markers.

Contextual factors may also have influenced participants’ expressions of their psychedelic experiences. The retreat setting, including group dynamics, interpersonal interactions, and heightened suggestibility [89,90] could amplify the phenomenon Durkheim described as “collective effervescence” [91,92], potentially shaping responses in ways unlikely to occur in controlled clinical settings.

Methodologically, the reliance on predefined word dictionaries—while a common analytical approach—may have missed nuances in slang, non-native speech, or culturally specific language [93]. Trained on Reddit comments, the automated emotion detection model may not fully capture the specialised language and emotional states associated with psychedelic integration, and imbalances in the training data [94] could bias the detection of underrepresented emotions. And crucially, the absence of a control group prevents us from ruling out alternative explanations for the observed changes, such as the known therapeutic benefits of journaling itself, which has been shown to improve mental health and well-being independent of any psychedelic intervention [95].

Finally, several data collection constraints should be noted. The voice journal format, though ecologically valid for capturing natural speech, may have led to self-censorship or altered expression due to participants’ awareness of being recorded. Inconsistencies in recording conditions - such as ambient noise, microphone distance, or device quality - could affect the reliability of acoustic data. Disruptions to participants’ schedules due to time zone differences, along with the absence of follow-up assessments, limit conclusions about the persistence of observed linguistic changes. Taken together, these factors suggest that the findings, while informative, should be interpreted with caution regarding their generalisability and precision.

### Future directions

This chatbot-based, speech-focused protocol exemplifies a scalable method for capturing nuanced psychological transformations around psychedelic sessions. By leveraging the widespread accessibility of smartphones and messaging applications, such automated approaches can reduce participant burden while enabling finer-grained tracking of integration processes in real-world settings. Future studies could expand this approach across diverse cultural contexts and larger sample sizes, or adapt it to other short-acting psychedelics (e.g., DMT) and longer-acting compounds (e.g., psilocybin, LSD). The inclusion of active control conditions—such as journaling without psychedelic use—or placebo arms will be essential for isolating the specific effects of psychedelic compounds from other therapeutic elements. Linking language or acoustic changes to neuroimaging, physiological, and behavioral measures could further elucidate the underlying mechanisms.

Longitudinal data collection at multiple post-dosing intervals (e.g., one and three months) will be critical for determining the persistence of observed linguistic and acoustic changes and their potential relation to lasting neuroplastic adaptations. Methodological refinements may include the use of advanced language models (e.g., GPT-4o, Llama 3, Claude 3.7) capable of detecting nuanced emotional tone, metaphorical richness, and non-standard linguistic constructions. However, the computational demands, “black box” nature, inherent biases, and ethical considerations associated with such models warrant careful attention. Transparent validation procedures, open-source dataset sharing, and interdisciplinary collaboration will be essential to ensuring that emerging computational tools contribute meaningfully and responsibly to both psychedelic science and clinical translation.

Finally, further investigation is needed into how different integration modalities (e.g., Bathje et al. 2022) shape post-session language trajectories, and whether personalised support can enhance therapeutic outcomes. These approaches should be tested across diverse populations, including clinical groups such as individuals with depression, who may engage with psychedelic experiences-and their integration-differently than healthy, experienced users. Ultimately, speech-based monitoring holds promise for guiding preparation, predicting outcomes, and supporting integration in both clinical and retreat settings.

### Conclusions

In conclusion, our data support the notion that everyday language and vocal patterns are sensitive indicators of psychedelic-induced psychological change. The observed shift toward introspective, cognitively oriented language-coupled with distinct vocal transitions-underscores how profoundly a single 5-MeO-DMT session can reshape one’s communication and sense of self. Not only do these methods offer new ways to map the trajectory of “preparation” and “integration,” but they also suggest a future where automated, real-time feedback might guide and support individuals undergoing psychedelic-assisted therapies or retreat experiences. By establishing the feasibility and utility of longitudinal voice analysis, our findings set the stage for broader efforts to illuminate the pathways through which psychedelics effect enduring personal growth and therapeutic benefits.

## Methods

### Ethical statement

This study adhered to the principles of the Helsinki Declaration and received approval from UCL’s Ethics Committee (ID: 19437/004). It was conducted in collaboration with the Tandava Retreat Centre (TRC), which supplied both facilitators and facilities. Participants reviewed the study details and provided informed consent online. Participation was voluntary, without compensation, and individuals could withdraw at any time without repercussion. All participants were provided with information about additional support resources in case of any study-related distress.

### Participants and study setting

Healthy, adult participants were recruited globally through the F.I.V.E. platform (www.five-meo.education), social media, and word of mouth. The final cohort (N = 29) was predominantly White/Caucasian (86.21%) and non-religious (93.10%), with prior experience with 5-MeO-DMT (mean lifetime use: 39 occasions (SD = 72.91, range = 1–300)).

### Procedures

#### Three-day retreat and 5-MeO-DMT session

All participants attended a three-day retreat at TRC. Day 1 focused on preparation, Day 2 on 5-MeO-DMT administration, and Day 3 on integration. On Day 2, participants vaporized a 12 mg dose of 5-MeO-DMT and inhaled it in a controlled setting under the supervision of experienced facilitators. For a comprehensive overview of study design and additional study components, such as EEG protocols, see Blackburne et al. (2024) [96].

#### Automated chatbot-based voice journal collection system

Voice journals were collected via *RetreatBot*, a custom-built chatbot hosted on the Telegram platform. Developed for Telegram using the OneReach.ai software platform Generative Studio X, this chatbot functioned as an EMA tool to capture voice journals of participants’ thoughts and feelings. Unlike more recent, dynamic conversational AIs (e.g., ChatGPT) that adapt in real-time, the chatbot employed in this study was rule-based, following a decision-tree model with predefined paths and functioning more like a structured app. A primary reason for choosing a chatbot was its inherently conversational interface (see: Fig. 1), which can reduce user burden and enhance engagement. In addition, the chatbot featured speech-based data entry-an approach suggested to be more intuitive and efficient than typing in mobile health applications [97]. By integrating these design elements, *RetreatBot* aimed to streamline participation and minimise the technological barriers or effort required from users.

Upon registration, each participant used a unique study ID, received an overview of the study procedures, and was guided on how to record voice journals. The default prompt was:

> *“Take a moment to reflect on your day and share your thoughts, feelings, and experiences. Record a voice note for about one minute.”*

Notifications were sent daily from seven days before (Day −7) to seven days after (Day +7) the dosing session. However, participants could also submit entries outside this core window to capture additional relevant experiences.

### Self-report measures

In this study, participants completed self-report instruments to assess psychedelic preparedness, subjective experiences (positive and negative), and post-experience well-being. We examined the relationships between these measures and participants’ voice journal data (both text- and audio-based features).

The **Psychedelic Preparedness Scale (PPS)** [23] includes 20 items rated on a seven-point Likert scale (1–7) designed to assess an individual’s readiness for a psychedelic experience. Each item is rated on a seven-point Likert scale (1–7), evaluating four key domains: preparatory readiness: knowledge-expectation, psychophysical readiness, intention-preparation, and support-planning. The total PPS score was calculated as the sum of all 20 items, resulting in a total range from 20 to 140.

The **Oceanic Boundlessness (OBN)** subscale of the 5-Dimensional Altered States of Consciousness Questionnaire (5D-ASC) [98,99] [98,99] includes 27 items rated on a visual analog scale from 0 (“Not at all”) to 100 (“Extremely”). It measures feelings of unity, spiritual experiences, blissful states, and insightfulness, capturing positively experienced depersonalization and derealization associated with heightened mood or euphoric exaltation. The OB score was calculated as the average of all OB-related items, providing a score in the 0–100 range.

The **Dread of Ego Dissolution (DED)** is another subscale of the 5D-ASC that assesses experiences related to ego disintegration, such as impaired control and cognition, anxiety, and disembodiment. It reflects negatively experienced derealisation and depersonalisation, including cognitive disturbances and loss of self-control. Scores for the DED subscale are calculated in the same manner as for OBN, yielding a score in the 0–100 range.

The **Emotional Breakthrough Inventory (EBI)** [79] includes 6 items rated on a six-point Likert scale (0–5), designed to assess the degree of emotional release or catharsis following a psychedelic experience. The total EBI score was calculated as the sum of all 6 items, yielding a score in the 0–30 range.

The **sWEMWBS** is a short version of the **Warwick–Edinburgh Mental Well-Being Scale** (WEMWBS) [100] and includes 7 of the WEMWBS’s 14 items rated on a five-point Likert scale (1–5), assessing both hedonic (e.g., happiness) and eudaimonic (e.g., purpose, connection) aspects of mental well-being. The total WEMWBS score was calculated as the sum of all 7 items, with a resulting range of 7 to 35.

### Data preprocessing and feature extraction

#### Voice journal preprocessing

All voice recordings were automatically transcribed upon submission using the AssemblyAI API [101]. Two researchers (AS and JK) subsequently performed manual checks to ensure transcription accuracy. Journal entries containing no spoken text were removed.

To maintain participant confidentiality, sensitive information (e.g., names, locations, organizations, dates) was anonymized. An automated approach using the transformers library’s TokenClassificationPipeline and a pre-trained named entity recognition model (Davlan/bert-base-multilingual-cased-ner-hrl) was used. Any remaining identifiable elements were further redacted manually.

#### Timestamp adjustments and relative dating

Universal Coordinated Time (UTC) timestamps were converted to the local retreat timezone (Mexico City). For consistency, the retreat start time was standardised to 8:00 AM. Each voice entry’s relative date was then determined as the difference (in days) between its local-adjusted timestamp and the retreat start. Entries were labeled “Pre” or “Post” depending on whether they occurred before or after the 5-MeO-DMT session (Day 0). Recordings made on the same calendar day as the dosing but referring to anticipated or immediate post-experience content were manually assigned slightly adjusted relative dates (−0.5 or +0.5) to ensure accurate pre–post categorisation. Voice entries from Day −14 through Day +14 were retained for analysis.

#### Text-based feature extraction

After transcription, each journal entry was split into sentences using the NLTK sentence tokenizer [102]. Two complementary approaches were used:

##### 1. Categorical Word Frequency (LIWC-22)

Each sentence was processed by Linguistic Inquiry and Word Count (LIWC-22) software (Pennebaker et al., 2022). LIWC scores represent the proportion of words within each text that fall into the specific category. For example, in the sentence *“I am happy,”* the score for *Affect* category would be 0.33, as one out of the three words (*“happy”*) pertains to affective content. Categories with an average frequency below 1% across all data were excluded to allow for reliable analysis of the coefficients [103]

##### 2. Emotion Classification (RoBERTa-GoEmotions)

A Robustly Optimised BERT Pretraining Approach (RoBERTa) model,, an optimised variant of the BERT model developed by Facebook AI [104], fine-tuned on the GoEmotions dataset (Demszky et al., 2020) was used to generate probability scores for 27 emotion categories plus a neutral class (<u>huggingface.co/SamLowe/roberta-base-go_emotions</u>). These emotions included positive (admiration, amusement, approval, caring, desire, excitement, gratitude, joy, love, optimism, pride, relief), negative (anger, annoyance, disappointment, disapproval, disgust, embarrassment, fear, grief, nervousness, remorse, sadness), and ambiguous (confusion, curiosity, realization, surprise) ones, following the GoEmotions taxonomy.

To represent these features for analysis, two main types of scores were computed:

● **Per-Sentence (PS) Scores:** Raw LIWC category or emotion scores calculated for each sentence.
● **Weighted Mean (WM) Scores:** Individual pre- and post- dosing averages weighted by sentence length (in words). WM was calculated using this formula:

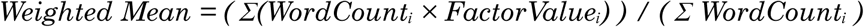

where *WordCount_i_* is the number of words in a given sentence, and *FactorValue_i_* represents the LIWC or emotional category score for that sentence. This formula calculates a weighted average by giving more influence to sentences with higher word counts, ensuring that longer sentences contribute proportionally to the overall score. By applying this weighting, we achieved a more representative measure of language patterns over the entire study period.

#### Acoustic features

Acoustic features were extracted from voice recordings using the OpenSMILE toolkit (eGeMAPSv02), producing 88 parameters (e.g., pitch, intensity, speech rate, voice quality, spectral measures). For each participant, we calculated simple mean values of key parameters in the two-week pre-session (Days −14 to 0) versus post-session (Days 0 to +14) windows.

### Data analysis

All statistical tests were two-tailed unless stated otherwise, with p-values adjusted via the Benjamini–Hochberg False Discovery Rate (FDR) method within each analysis family (e.g., LIWC features, emotion features, acoustic features). We used R²-adjusted and the Bayesian Information Criterion (BIC) to assess regression model performance and guide model selection.

#### Pre– to post–dosing comparisons

##### Text-based features

Linear mixed-effects models were employed to examine changes in LIWC and emotion category usage from pre- to post-session:

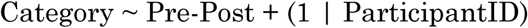

Here, **Category** represents a LIWC or RoBERTa-GoEmotions measure, **Pre-Post** is a fixed effect, and **ParticipantID** is a random intercept to account for repeated measures. Statistically significant shifts were identified based on FDR-corrected p-values.

##### Acoustic metrics

Paired t-tests were used to compare pre- vs. post-dosing means of each acoustic feature. Differences were standardized as z-scores, and 95% confidence intervals were computed for each parameter.

#### Temporal analyses

We conducted a day-by-day examination of how certain linguistic and vocal features evolved from Day −14 to Day +14. For linguistic metrics (i.e., “Cognition,” and “Social” words), we calculated participant-level z-scores relative to each individual’s overall mean, then aggregated these daily across the entire sample. Smoothing with a Gaussian kernel of 1.5 days was applied for visualisation. To facilitate discrete comparisons, the 28-day window was divided into four periods: Week −2 (Days −14 to −7), Week −1 (Days y7 to 0), Week +1 (Days 0 to 7), and Week +2 (Days 7 to 14). Mean standardized usage of linguistic categories in each period was compared using pairwise t-tests.

Similarly, for acoustic characteristics (pitch, jitter, shimmer), baseline values for all participants were calculated as the mean and standard deviation (±1 SD) across all available recordings. Deviations beyond this range were classified as “High” or “Low” deviations for each acoustic feature. These deviations were calculated separately for pitch (in Hz, derived from semitone values relative to 27.5 Hz), jitter (local), and shimmer (local dB). Deviation frequencies were aggregated at the daily level, then smoothed using a Gaussian kernel (σ = 1). Group-level comparisons were performed across four time periods—Week −2 (Days −14 to −7), Week −1 (Days −7 to 0), Week +1 (Days 0 to 7), and Week +2 (Days 7 to 14)—using independent two-sample t-tests. False discovery rate (FDR) correction was applied across all pairwise tests. Deviations were visualized as daily time series and as period-averaged bar plots, annotated with statistical significance where applicable.

#### Predictive modeling

To explore whether pre-session linguistic or acoustic patterns predicted subjective experiences, we first conducted a correlation-based feature selection (|r| > 0.6) between pre-dosing weighted means of LIWC categories and self-report measures (PPS, OBN, DED, EBI, sWEMWBS). Retained features were then entered into linear regressions as predictors of these psychometric outcomes. To evaluate model robustness and capture multivariate effects across the full feature set, ridge regression was additionally performed using all pre-dosing weighted means of LIWC features as predictors. Ridge models were optimized via 10-fold cross-validation to select the regularization parameter (alpha), and the top five standardized coefficients were extracted for interpretability.

For the emotion-based features, we used Principal Component Analysis (PCA) on weighted mean pre-dosing RoBERTa-GoEmotions scores. Components were selected based on an elbow plot, and emotions with loadings ≥ |0.2| were used to interpret each component. Regression analyses tested whether these component scores predicted the five self-report measures. Statistically significant models were identified through adjusted R², BIC, and p-values.

## Acknowledgements

We wish to thank Tandava Retreats for their generous assistance with the data collection process, with special recognition to Victoria Wueschner and Joel Brierre for their key contributions in facilitating the 5-MeO-DMT sessions. We are also grateful to Luis Fabian Rodriguez, Otto Maier, James Sanders, and George Deane for their support during data collection. Our deepest appreciation goes to the participants who volunteered for this study, generously contributing their time and personal voice journals. This research wouldn’t be possible without their openness. Additionally, we thank the contributors to the crowdfunding campaign, whose support made this research possible.

## Funding statement

J.K. is supported by UCL’s Computing Career Development Studentship. The development of experience sampling chatbot for the study was supported by an academic fellowship received by J.K. from the technology company OneReach.ai, which provided the access to *Generative Studio X* allowing for building and deploying the chatbot, associated training, and technical support. OneReach.ai had no role in the study’s design, data collection, analysis, or interpretation. G.B. receives support from the Leverhulme Trust, R.G.M. is funded by the Wellcome Trust, and J.I.S. is supported by Wellcome Leap.

## Competing interests statement

JK is currently an employee at Compass Pathways plc. This work is conducted independently and not associated with Compass Pathways plc. Other authors have no conflicts of interest.

## Authors’ contributions

**J.K.:** Conceptualisation, Methodology, Software, Formal analysis, Data curation, Visualisation, Writing – original draft, Funding acquisition, Project administration. **R.M.:** Conceptualisation, Investigation, Methodology, Writing – review & editing. **A.S.:** Data curation (qualitative), Validation. **G.B.:** Project administration, Investigation, Writing - Review & Editing. **D.L.:** Software, Writing – review & editing. **J.S.:** Supervision, Conceptualisation, Project administration, Writing – review & editing.

## Notes

### Summary of Updates

Included the acknowledgements section, which was unintentionally omitted from the previous version.

