## Supplementary_Material for "Speech markers of psychedelic-induced psychological change"

| Measure | Pre-dosing | Post-dosing |
|---|---|---|
| Age | 48.1 ± 10.9<br>(n=26) | - |
| Gender | Male: 15 (57.7%);<br>Female: 11 (42.3%) | - |
| (DASS) Total | 27.27 ± 4.07 | 24.65 ± 4.88 |
| (DASS) Depression | 16.08 ± 2.30 | 15.62 ± 1.96 |
| (DASS) Anxiety | 17.00 ± 2.73 | 15.00 ± 2.35 |
| (DASS) Stress | 21.46 ± 5.76 | 18.69 ± 6.88 |
| (sWEMWBS) Wellbeing | 26.92 ± 4.52 | 29.69 ± 3.25 |
| (EBI) Emotional Breakthrough | - | 257.35 ± 186.33 |
| (ASC) Oceanic Boundlessness | - | 0.71 ± 0.27 |
| (ASC) Dread of Ego Dissolution | - | 0.28 ± 0.27 |

**Supplementary Table S1**
*Demographic characteristics and clinical measures before and after 5-MeO-DMT administration.*
The table reports mean ± standard deviation for continuous variables and counts (percentages) for categorical variables. Pre-dosing measures were collected 2 weeks before the ceremony, while post-dosing measures were collected 2 weeks after, except for acute effects measures (EBI, ASC) which were collected 1 day post-dosing. Data are shown for participants who completed all psychometric questionnaires (n=26) out of the total 29 participants who provided voice recordings. DASS subscales measure depression, anxiety, and stress symptoms, with higher scores indicating greater symptom severity. sWEMWBS measures psychological wellbeing, with higher scores indicating greater wellbeing. EBI measures emotional breakthrough experiences, ASC-OBN measures oceanic boundlessness experiences, and ASC-DED measures dread of ego dissolution during the acute psychedelic state.

| Main Category | Subcategory | Raw Category | Coefficient | Std. Error | CI Lower | CI Upper | P-Val | FDR Corr. P-Val | N Observ. | Sig. |
|---|---|---|---|---|---|---|---|---|---|---|
| Affect | Affect (Total) | Affect | -0.20 | 0.57 | -1.32 | 0.91 | 0.720 | 0.855 | 4127 | ns |
| Affect | Positive Emotion | emo_pos | 0.10 | 0.27 | -0.42 | 0.63 | 0.707 | 0.855 | 4127 | ns |
| Affect | Emotion | emotion | -0.22 | 0.30 | -0.80 | 0.36 | 0.453 | 0.727 | 4127 | ns |
| Affect | Positive Tone | tone_pos | 0.17 | 0.56 | -0.93 | 1.27 | 0.760 | 0.855 | 4127 | ns |
| Cognition | Cognition (Total) | Cognition | 1.96 | 0.46 | 1.06 | 2.86 | 0.000 | 0.000 | 4127 | *** |
| Cognition | All-or-none | allnone | -0.06 | 0.19 | -0.42 | 0.30 | 0.741 | 0.855 | 4127 | ns |
| Cognition | Certitude | certitude | 0.43 | 0.16 | 0.12 | 0.75 | 0.007 | 0.037 | 4127 | * |
| Cognition | Cognitive Processes | cogproc | 1.93 | 0.43 | 1.08 | 2.77 | 0.000 | 0.000 | 4127 | *** |
| Cognition | Differentiation | differ | 0.44 | 0.21 | 0.03 | 0.85 | 0.034 | 0.096 | 4127 | ns |
| Cognition | Insight | insight | 0.63 | 0.23 | 0.18 | 1.07 | 0.006 | 0.037 | 4127 | * |
| Cognition | Tentative | tentat | 0.74 | 0.21 | 0.33 | 1.16 | 0.000 | 0.005 | 4127 | ** |
| Conversation | Conversation (Total) | Conversation | -1.64 | 0.58 | -2.78 | -0.51 | 0.005 | 0.034 | 4127 | * |
| Conversation | Assent | assent | -1.25 | 0.52 | -2.28 | -0.22 | 0.017 | 0.059 | 4127 | ns |
| Drives | Drives (Total) | Drives | -0.01 | 0.19 | -0.38 | 0.37 | 0.966 | 0.966 | 4127 | ns |
| Drives | Affiliation | affiliation | -0.16 | 0.15 | -0.46 | 0.14 | 0.297 | 0.534 | 4127 | ns |
| Lifestyle | Lifestyle (Total) | Lifestyle | -0.10 | 0.20 | -0.49 | 0.30 | 0.626 | 0.854 | 4127 | ns |
| Linguistic | Adjectives | adj | 0.49 | 0.39 | -0.28 | 1.26 | 0.210 | 0.395 | 4127 | ns |
| Linguistic | Adverbs | adverb | 1.04 | 0.39 | 0.28 | 1.81 | 0.008 | 0.037 | 4127 | * |
| Linguistic | Articles | article | -0.02 | 0.21 | -0.44 | 0.40 | 0.931 | 0.966 | 4127 | ns |
| Linguistic | Auxiliary Verbs | auxverb | 0.80 | 0.34 | 0.12 | 1.47 | 0.020 | 0.066 | 4127 | ns |
| Linguistic | Conjunctions | conj | -0.21 | 0.32 | -0.83 | 0.42 | 0.512 | 0.750 | 4127 | ns |
| Linguistic | Determiners | det | -0.23 | 0.35 | -0.92 | 0.46 | 0.517 | 0.750 | 4127 | ns |
| Linguistic | Negations | negate | 0.21 | 0.14 | -0.06 | 0.48 | 0.122 | 0.282 | 4127 | ns |
| Linguistic | Numbers | number | 0.04 | 0.13 | -0.21 | 0.30 | 0.754 | 0.855 | 4127 | ns |
| Linguistic | Prepositions | prep | -0.26 | 0.34 | -0.93 | 0.40 | 0.441 | 0.727 | 4127 | ns |
| Linguistic | Quantities | quantity | -0.37 | 0.30 | -0.95 | 0.21 | 0.211 | 0.395 | 4127 | ns |
| Linguistic | Verbs | verb | 0.67 | 0.49 | -0.28 | 1.62 | 0.168 | 0.343 | 4127 | ns |
| Motives | Allure | allure | 0.83 | 0.50 | -0.15 | 1.81 | 0.096 | 0.241 | 4127 | ns |
| Perception | Perception (Total) | Perception | 0.04 | 0.40 | -0.73 | 0.82 | 0.911 | 0.966 | 4127 | ns |
| Perception | Feeling | feeling | 0.43 | 0.19 | 0.05 | 0.81 | 0.026 | 0.079 | 4127 | ns |
| Perception | Motion | motion | -0.14 | 0.15 | -0.43 | 0.16 | 0.359 | 0.621 | 4127 | ns |
| Perception | Space | space | 0.17 | 0.29 | -0.39 | 0.73 | 0.548 | 0.771 | 4127 | ns |
| Physical | Physical (Total) | Physical | 0.08 | 0.22 | -0.35 | 0.51 | 0.721 | 0.855 | 4127 | ns |
| Pronoun | 1st Person Singular | i | 0.40 | 0.28 | -0.15 | 0.94 | 0.156 | 0.335 | 4127 | ns |
| Pronoun | Impersonal Pronouns | ipron | 0.13 | 0.35 | -0.54 | 0.81 | 0.699 | 0.855 | 4127 | ns |
| Pronoun | Personal Pronouns | ppron | -0.02 | 0.33 | -0.68 | 0.63 | 0.948 | 0.966 | 4127 | ns |
| Social | Social (Total) | Social | -1.69 | 0.64 | -2.93 | -0.44 | 0.008 | 0.037 | 4127 | * |
| Social | Communication | comm | -1.22 | 0.49 | -2.17 | -0.26 | 0.013 | 0.047 | 4127 | * |
| Social | Politeness | polite | -0.78 | 0.51 | -1.78 | 0.22 | 0.126 | 0.282 | 4127 | ns |
| Social | Prosocial Behavior | prosocial | -0.16 | 0.24 | -0.63 | 0.31 | 0.512 | 0.750 | 4127 | ns |
| Social | Social Behavior | socbehav | -1.02 | 0.57 | -2.14 | 0.09 | 0.073 | 0.193 | 4127 | ns |
| Social | Social Referents | socrefs | -0.64 | 0.25 | -1.14 | -0.14 | 0.012 | 0.047 | 4127 | * |
| Time | Future Focus | focusfuture | -0.58 | 0.18 | -0.93 | -0.24 | 0.001 | 0.008 | 4127 | ** |
| Time | Past Focus | focuspast | 1.60 | 0.27 | 1.07 | 2.12 | 0.000 | 0.000 | 4127 | *** |
| Time | Present Focus | focuspresent | -0.04 | 0.32 | -0.68 | 0.60 | 0.902 | 0.966 | 4127 | ns |

**Supplementary Table S2**

*Mixed-effects model results for changes in LIWC vocabulary category usage in participants' voice journals following 5-MeO-DMT administration.*

The table reports coefficients, standard errors, 95% confidence intervals, uncorrected p-values, and FDR-corrected p-values (Benjamini-Hochberg) for each LIWC category and subcategory. Estimates reflect the average change in word category usage from pre- to post-dosing (n = 4127 sentences from 29 participants), based on mixed-effects models with a binary "Pre-Post" predictor and a random intercept for participant (model: PrePost + (1 | ParticipantID)). The Sig. column indicates the statistical significance of FDR-corrected p-values using the following notation: $p < 0.05$ (*), $p < 0.01$ (**), $p < 0.001$ (***), and ns for non-significant results.

| Category | Coefficient | Standard Error | CI Lower | CI Upper | P-Value | Corrected P-Value | N Observ. | Sig. |
|---|---|---|---|---|---|---|---|---|
| admiration | 0.0278 | 0.01 | 0.01 | 0.04 | 0.000 | 0.002 | 4127 | ** |
| joy | 0.0217 | 0.01 | 0.01 | 0.04 | 0.002 | 0.011 | 4127 | * |
| realization | 0.0087 | 0.00 | 0.00 | 0.01 | 0.004 | 0.014 | 4127 | * |
| love | 0.0051 | 0.00 | 0.00 | 0.01 | 0.122 | 0.284 | 4127 | ns |
| approval | 0.0051 | 0.01 | -0.01 | 0.02 | 0.334 | 0.491 | 4127 | ns |
| sadness | 0.0046 | 0.00 | 0.00 | 0.01 | 0.176 | 0.306 | 4127 | ns |
| gratitude | 0.003 | 0.00 | -0.01 | 0.01 | 0.532 | 0.621 | 4127 | ns |
| confusion | 0.0028 | 0.00 | 0.00 | 0.01 | 0.467 | 0.594 | 4127 | ns |
| relief | 0.0027 | 0.00 | 0.00 | 0.00 | 0.000 | 0.002 | 4127 | ** |
| surprise | 0.0024 | 0.00 | 0.00 | 0.01 | 0.186 | 0.306 | 4127 | ns |
| remorse | 0.0021 | 0.00 | 0.00 | 0.01 | 0.147 | 0.294 | 4127 | ns |
| amusement | 0.0011 | 0.00 | 0.00 | 0.00 | 0.463 | 0.594 | 4127 | ns |
| embarrassment | 0.001 | 0.00 | 0.00 | 0.00 | 0.141 | 0.294 | 4127 | ns |
| pride | 0.0007 | 0.00 | 0.00 | 0.00 | 0.025 | 0.077 | 4127 | ns |
| disgust | 0.0004 | 0.00 | 0.00 | 0.00 | 0.595 | 0.666 | 4127 | ns |
| grief | 0.0001 | 0.00 | 0.00 | 0.00 | 0.294 | 0.458 | 4127 | ns |
| anger | -0.0003 | 0.00 | 0.00 | 0.00 | 0.720 | 0.720 | 4127 | ns |
| disapproval | -0.0011 | 0.00 | -0.01 | 0.00 | 0.665 | 0.716 | 4127 | ns |
| disappointment | -0.0011 | 0.00 | -0.01 | 0.00 | 0.694 | 0.720 | 4127 | ns |
| annoyance | -0.0016 | 0.00 | -0.01 | 0.00 | 0.383 | 0.536 | 4127 | ns |
| desire | -0.0016 | 0.00 | -0.01 | 0.00 | 0.522 | 0.621 | 4127 | ns |
| fear | -0.0041 | 0.00 | -0.01 | 0.00 | 0.003 | 0.014 | 4127 | * |
| curiosity | -0.005 | 0.00 | -0.01 | 0.00 | 0.055 | 0.139 | 4127 | ns |
| optimism | -0.0051 | 0.00 | -0.01 | 0.00 | 0.172 | 0.306 | 4127 | ns |
| caring | -0.0058 | 0.00 | -0.01 | 0.00 | 0.019 | 0.067 | 4127 | ns |
| nervousness | -0.0066 | 0.00 | -0.01 | 0.00 | 0.000 | 0.002 | 4127 | ** |
| excitement | -0.022 | 0.00 | -0.03 | -0.01 | 0.000 | 0.000 | 4127 | *** |
| neutral | -0.0225 | 0.01 | -0.05 | 0.00 | 0.054 | 0.139 | 4127 | ns |

**Supplementary Table S3**

*Mixed-effects model results for changes in emotion expression in participants' voice journals following 5-MeO-DMT administration, as identified using a RoBERTa-based GoEmotions classifier.*

The table reports coefficients, standard errors, 95% confidence intervals, uncorrected p-values, and FDR-corrected p-values (Benjamini-Hochberg) for each emotion category. Estimates reflect average changes in model-predicted emotion probabilities from pre- to post-dosing (n = 4127 sentences from 29 participants), using mixed-effects models with a binary "Pre-Post" predictor and random intercept for participant (model: PrePost + (1 | ParticipantID)). The Sig. column indicates the statistical significance of FDR-corrected p-values using the following notation: $p < 0.05$ (*), $p < 0.01$ (**), $p < 0.001$ (***), and ns for non-significant results.

| Feature | Pre Mean | Post Mean | Mean Difference | Standard Error | CI Lower | CI Upper | P-Value | FDR Corr. P-Val | N Observ. | Sig. |
|---|---|---|---|---|---|---|---|---|---|---|
| jitterLocal_sma3nz_stddevNorm | 1.99 | 1.84 | -0.91 | 0.21 | -1.32 | -0.51 | 0 | 0.021 | 23 | * |
| shimmerLocaldB_sma3nz_amean | 1.25 | 1.34 | 0.76 | 0.21 | 0.35 | 1.17 | 0.001 | 0.045 | 23 | * |
| jitterLocal_sma3nz_amean | 0.05 | 0.05 | 0.75 | 0.21 | 0.35 | 1.16 | 0.002 | 0.045 | 23 | * |
| VoicedSegmentsPerSec | 2.17 | 2.41 | 0.69 | 0.21 | 0.28 | 1.10 | 0.003 | 0.055 | 23 | ns |
| F0semitoneFrom27.5Hz_sma3nz_percentile50.0 | 25.41 | 24.02 | -0.68 | 0.21 | -1.09 | -0.27 | 0.004 | 0.055 | 23 | ns |
| HNRdBACF_sma3nz_amean | 3.17 | 2.22 | -0.68 | 0.21 | -1.08 | -0.27 | 0.004 | 0.055 | 23 | ns |
| logRelF0-H1-A3_sma3nz_stddevNorm | 0.71 | 0.85 | 0.62 | 0.21 | 0.21 | 1.03 | 0.007 | 0.085 | 23 | ns |
| F0semitoneFrom27.5Hz_sma3nz_percentile80.0 | 28.27 | 27.35 | -0.53 | 0.21 | -0.94 | -0.12 | 0.018 | 0.167 | 23 | ns |
| MeanVoicedSegmentLengthSec | 0.31 | 0.28 | -0.53 | 0.21 | -0.94 | -0.12 | 0.018 | 0.167 | 23 | ns |
| F0semitoneFrom27.5Hz_sma3nz_amean | 25.03 | 23.91 | -0.53 | 0.21 | -0.94 | -0.12 | 0.019 | 0.167 | 23 | ns |
| spectralFlux_sma3_amean | 0.43 | 0.53 | 0.45 | 0.21 | 0.05 | 0.86 | 0.04 | 0.28 | 23 | ns |
| spectralFluxUV_sma3nz_amean | 0.23 | 0.33 | 0.45 | 0.21 | 0.04 | 0.86 | 0.041 | 0.28 | 23 | ns |
| logRelF0-H1-A3_sma3nz_amean | 23.97 | 22.59 | -0.45 | 0.21 | -0.86 | -0.04 | 0.041 | 0.28 | 23 | ns |
| slopeV0-500_sma3nz_amean | 0.02 | 0.01 | -0.43 | 0.21 | -0.84 | -0.02 | 0.051 | 0.313 | 23 | ns |
| shimmerLocaldB_sma3nz_stddevNorm | 0.85 | 0.82 | -0.42 | 0.21 | -0.83 | -0.02 | 0.054 | 0.313 | 23 | ns |
| spectralFluxV_sma3nz_amean | 0.54 | 0.64 | 0.42 | 0.21 | 0.01 | 0.83 | 0.057 | 0.313 | 23 | ns |
| F0semitoneFrom27.5Hz_sma3nz_meanRisingSlope | 330.51 | 303.83 | -0.38 | 0.21 | -0.79 | 0.03 | 0.081 | 0.42 | 23 | ns |
| slopeUV0-500_sma3nz_amean | 0.01 | 0.00 | -0.37 | 0.21 | -0.78 | 0.03 | 0.086 | 0.422 | 23 | ns |
| F0semitoneFrom27.5Hz_sma3nz_percentile20.0 | 20.06 | 18.88 | -0.36 | 0.21 | -0.77 | 0.05 | 0.101 | 0.466 | 23 | ns |
| MeanUnvoicedSegmentLength | 0.19 | 0.15 | -0.32 | 0.21 | -0.72 | 0.09 | 0.145 | 0.638 | 23 | ns |
| slopeV0-500_sma3nz_stddevNorm | 4.11 | -6.33 | -0.30 | 0.21 | -0.71 | 0.11 | 0.161 | 0.675 | 23 | ns |
| mfcc3_sma3_amean | 11.77 | 12.82 | 0.29 | 0.21 | -0.12 | 0.70 | 0.174 | 0.696 | 23 | ns |
| mfcc3V_sma3nz_stddevNorm | 1.32 | 1.78 | 0.27 | 0.21 | -0.13 | 0.68 | 0.202 | 0.743 | 23 | ns |
| logRelF0-H1-H2_sma3nz_stddevNorm | 13.98 | -17.70 | -0.27 | 0.21 | -0.68 | 0.14 | 0.213 | 0.743 | 23 | ns |
| loudness_sma3_pctlrange0-2 | 0.83 | 0.88 | 0.24 | 0.21 | -0.17 | 0.65 | 0.255 | 0.743 | 23 | ns |
| loudness_sma3_meanFallingSlope | 8.00 | 8.51 | 0.24 | 0.21 | -0.17 | 0.64 | 0.271 | 0.743 | 23 | ns |
| mfcc2_sma3_stddevNorm | 20.49 | -1.23 | -0.24 | 0.21 | -0.64 | 0.17 | 0.272 | 0.743 | 23 | ns |
| slopeV500-1500_sma3nz_stddevNorm | -1.86 | -0.90 | 0.23 | 0.21 | -0.18 | 0.64 | 0.277 | 0.743 | 23 | ns |
| F0semitoneFrom27.5Hz_sma3nz_stddevRisingSlope | 479.93 | 449.01 | -0.23 | 0.21 | -0.64 | 0.18 | 0.283 | 0.743 | 23 | ns |
| StddevUnvoicedSegmentLength | 0.26 | 0.21 | -0.23 | 0.21 | -0.64 | 0.18 | 0.283 | 0.743 | 23 | ns |
| loudness_sma3_percentile80.0 | 1.06 | 1.12 | 0.22 | 0.21 | -0.19 | 0.63 | 0.295 | 0.743 | 23 | ns |
| loudness_sma3_stddevRisingSlope | 7.88 | 8.29 | 0.22 | 0.21 | -0.19 | 0.63 | 0.305 | 0.743 | 23 | ns |
| loudness_sma3_meanRisingSlope | 11.31 | 11.85 | 0.21 | 0.21 | -0.20 | 0.62 | 0.321 | 0.743 | 23 | ns |
| F3amplitudeLogRelF0_sma3nz_amean | -89.06 | -85.51 | 0.20 | 0.21 | -0.21 | 0.61 | 0.351 | 0.743 | 23 | ns |
| mfcc4_sma3_amean | 1.36 | 2.40 | 0.20 | 0.21 | -0.21 | 0.61 | 0.358 | 0.743 | 23 | ns |
| mfcc3V_sma3nz_amean | 13.57 | 14.32 | 0.19 | 0.21 | -0.22 | 0.60 | 0.363 | 0.743 | 23 | ns |
| loudness_sma3_amean | 0.68 | 0.72 | 0.19 | 0.21 | -0.22 | 0.60 | 0.379 | 0.743 | 23 | ns |
| loudness_sma3_percentile50.0 | 0.56 | 0.60 | 0.19 | 0.21 | -0.22 | 0.59 | 0.383 | 0.743 | 23 | ns |
| StddevVoicedSegmentLengthSec | 0.32 | 0.31 | -0.19 | 0.21 | -0.59 | 0.22 | 0.383 | 0.743 | 23 | ns |
| mfcc4_sma3_stddevNorm | -4.66 | -51.87 | -0.18 | 0.21 | -0.59 | 0.23 | 0.39 | 0.743 | 23 | ns |
| alphaRatioV_sma3nz_stddevNorm | -0.65 | -1.42 | -0.18 | 0.21 | -0.59 | 0.23 | 0.391 | 0.743 | 23 | ns |
| F2amplitudeLogRelF0_sma3nz_amean | -85.26 | -81.88 | 0.18 | 0.21 | -0.23 | 0.59 | 0.401 | 0.743 | 23 | ns |
| F1amplitudeLogRelF0_sma3nz_amean | -79.02 | -75.38 | 0.18 | 0.21 | -0.23 | 0.59 | 0.402 | 0.743 | 23 | ns |
| logRelF0-H1-H2_sma3nz_amean | 4.86 | 4.38 | -0.18 | 0.21 | -0.59 | 0.23 | 0.403 | 0.743 | 23 | ns |
| loudness_sma3_stddevFallingSlope | 6.29 | 6.58 | 0.17 | 0.21 | -0.24 | 0.58 | 0.413 | 0.743 | 23 | ns |
| mfcc4V_sma3nz_amean | 1.00 | 2.05 | 0.17 | 0.21 | -0.24 | 0.58 | 0.415 | 0.743 | 23 | ns |
| alphaRatioUV_sma3nz_amean | -6.90 | -7.39 | -0.17 | 0.21 | -0.58 | 0.24 | 0.428 | 0.743 | 23 | ns |
| F3bandwidth_sma3nz_amean | 844.13 | 834.88 | -0.17 | 0.21 | -0.58 | 0.24 | 0.432 | 0.743 | 23 | ns |
| slopeV500-1500_sma3nz_amean | -0.02 | -0.02 | 0.17 | 0.21 | -0.24 | 0.57 | 0.438 | 0.743 | 23 | ns |
| mfcc4V_sma3nz_stddevNorm | -5.47 | -1.36 | 0.16 | 0.21 | -0.25 | 0.57 | 0.441 | 0.743 | 23 | ns |
| spectralFlux_sma3_stddevNorm | 1.10 | 1.07 | -0.16 | 0.21 | -0.57 | 0.25 | 0.446 | 0.743 | 23 | ns |
| hammarbergIndexUV_sma3nz_amean | 15.14 | 15.66 | 0.16 | 0.21 | -0.25 | 0.57 | 0.451 | 0.743 | 23 | ns |
| HNRdBACF_sma3nz_stddevNorm | 1.36 | -2.47 | -0.16 | 0.21 | -0.57 | 0.25 | 0.455 | 0.743 | 23 | ns |
| loudness_sma3_percentile20.0 | 0.23 | 0.25 | 0.16 | 0.21 | -0.25 | 0.57 | 0.459 | 0.743 | 23 | ns |
| loudnessPeaksPerSec | 2.49 | 2.53 | 0.16 | 0.21 | -0.25 | 0.56 | 0.465 | 0.743 | 23 | ns |
| equivalentSoundLevel_dBp | -28.55 | -28.13 | 0.15 | 0.21 | -0.26 | 0.56 | 0.488 | 0.766 | 23 | ns |
| loudness_sma3_stddevNorm | 0.77 | 0.76 | -0.14 | 0.21 | -0.55 | 0.27 | 0.514 | 0.787 | 23 | ns |
| mfcc3_sma3_stddevNorm | 1.20 | 1.52 | 0.13 | 0.21 | -0.28 | 0.54 | 0.531 | 0.787 | 23 | ns |
| F0semitoneFrom27.5Hz_sma3nz_stddevNorm | 0.26 | 0.26 | 0.13 | 0.21 | -0.28 | 0.54 | 0.536 | 0.787 | 23 | ns |
| mfcc1V_sma3nz_amean | 31.46 | 31.09 | -0.13 | 0.21 | -0.54 | 0.28 | 0.536 | 0.787 | 23 | ns |
| F2frequency_sma3nz_stddevNorm | 0.19 | 0.18 | -0.13 | 0.21 | -0.53 | 0.28 | 0.556 | 0.802 | 23 | ns |
| F3frequency_sma3nz_stddevNorm | 0.12 | 0.12 | 0.12 | 0.21 | -0.29 | 0.52 | 0.588 | 0.834 | 23 | ns |

| | | | | | | | | | | |
|---|---|---|---|---|---|---|---|---|---|---|
| hammarbergIndexV_sma3nz_amean | 27.36 | 27.03 | -0.11 | 0.21 | -0.52 | 0.30 | 0.615 | 0.859 | 23 | ns |
| F3frequency_sma3nz_amean | 2683.28 | 2677.74 | -0.10 | 0.21 | -0.51 | 0.31 | 0.641 | 0.873 | 23 | ns |
| F1frequency_sma3nz_stddevNorm | 0.40 | 0.40 | 0.10 | 0.21 | -0.31 | 0.51 | 0.646 | 0.873 | 23 | ns |
| F0semitoneFrom27.5Hz_sma3nz_pctlrange0-2 | 8.21 | 8.48 | 0.09 | 0.21 | -0.31 | 0.50 | 0.655 | 0.873 | 23 | ns |
| slopeUV500-1500_sma3nz_amean | -0.01 | -0.01 | -0.09 | 0.21 | -0.50 | 0.32 | 0.683 | 0.897 | 23 | ns |
| F1bandwidth_sma3nz_amean | 1235.21 | 1230.58 | -0.08 | 0.21 | -0.49 | 0.33 | 0.712 | 0.922 | 23 | ns |
| spectralFluxV_sma3nz_stddevNorm | 0.86 | 0.87 | 0.07 | 0.21 | -0.33 | 0.48 | 0.725 | 0.925 | 23 | ns |
| F0semitoneFrom27.5Hz_sma3nz_stddevFallingSlope | 246.61 | 239.49 | -0.07 | 0.21 | -0.48 | 0.34 | 0.743 | 0.934 | 23 | ns |
| F2frequency_sma3nz_amean | 1605.20 | 1601.85 | -0.06 | 0.21 | -0.47 | 0.35 | 0.773 | 0.955 | 23 | ns |
| mfcc2V_sma3nz_stddevNorm | -0.85 | -0.17 | 0.06 | 0.21 | -0.35 | 0.47 | 0.788 | 0.955 | 23 | ns |
| F3amplitudeLogRelF0_sma3nz_stddevNorm | -0.88 | -0.88 | -0.06 | 0.21 | -0.46 | 0.35 | 0.793 | 0.955 | 23 | ns |
| mfcc2_sma3_amean | 9.08 | 9.31 | 0.05 | 0.21 | -0.36 | 0.46 | 0.809 | 0.962 | 23 | ns |
| F2bandwidth_sma3nz_amean | 932.82 | 929.78 | -0.05 | 0.21 | -0.46 | 0.36 | 0.828 | 0.97 | 23 | ns |
| F1amplitudeLogRelF0_sma3nz_stddevNorm | -1.15 | -1.16 | -0.04 | 0.21 | -0.45 | 0.37 | 0.847 | 0.97 | 23 | ns |
| F3bandwidth_sma3nz_stddevNorm | 0.49 | 0.49 | 0.04 | 0.21 | -0.37 | 0.45 | 0.849 | 0.97 | 23 | ns |
| mfcc1_sma3_amean | 24.45 | 24.57 | 0.04 | 0.21 | -0.37 | 0.44 | 0.868 | 0.979 | 23 | ns |
| F1bandwidth_sma3nz_stddevNorm | 0.21 | 0.21 | 0.02 | 0.21 | -0.39 | 0.43 | 0.913 | 0.992 | 23 | ns |
| F2bandwidth_sma3nz_stddevNorm | 0.40 | 0.40 | 0.02 | 0.21 | -0.39 | 0.43 | 0.935 | 0.992 | 23 | ns |
| alphaRatioV_sma3nz_amean | -16.91 | -16.88 | 0.01 | 0.21 | -0.40 | 0.42 | 0.95 | 0.992 | 23 | ns |
| F1frequency_sma3nz_amean | 567.34 | 567.72 | 0.01 | 0.21 | -0.40 | 0.42 | 0.964 | 0.992 | 23 | ns |
| mfcc2V_sma3nz_amean | 10.26 | 10.22 | -0.01 | 0.21 | -0.42 | 0.40 | 0.968 | 0.992 | 23 | ns |
| F2amplitudeLogRelF0_sma3nz_stddevNorm | -0.96 | -0.96 | 0.01 | 0.21 | -0.40 | 0.42 | 0.975 | 0.992 | 23 | ns |
| mfcc1_sma3_stddevNorm | 0.68 | 0.68 | 0.01 | 0.21 | -0.40 | 0.41 | 0.979 | 0.992 | 23 | ns |
| hammarbergIndexV_sma3nz_stddevNorm | 0.40 | 0.40 | 0.00 | 0.21 | -0.41 | 0.41 | 0.986 | 0.992 | 23 | ns |
| F0semitoneFrom27.5Hz_sma3nz_meanFallingSlope | 132.90 | 132.74 | 0.00 | 0.21 | -0.41 | 0.41 | 0.987 | 0.992 | 23 | ns |
| mfcc1V_sma3nz_stddevNorm | 0.41 | 0.41 | 0.00 | 0.21 | -0.41 | 0.41 | 0.992 | 0.992 | 23 | ns |

**Supplementary Table S4**

*Paired-sample t-test results for changes in 88 acoustic features extracted from participants' voice recordings before and after 5-MeO-DMT administration, using the openSMILE (eGeMAPSv02) toolkit (n = 23 participants).*

The table includes pre- and post-dosing means, mean differences, standard errors, 95% confidence intervals, uncorrected and FDR-corrected p-values (Benjamini-Hochberg), and significance levels. The Sig. column indicates the statistical significance of FDR-corrected p-values using the following notation: $p < 0.05$ (), $p < 0.01$ (), $p < 0.001$ (), and ns for non-significant results.

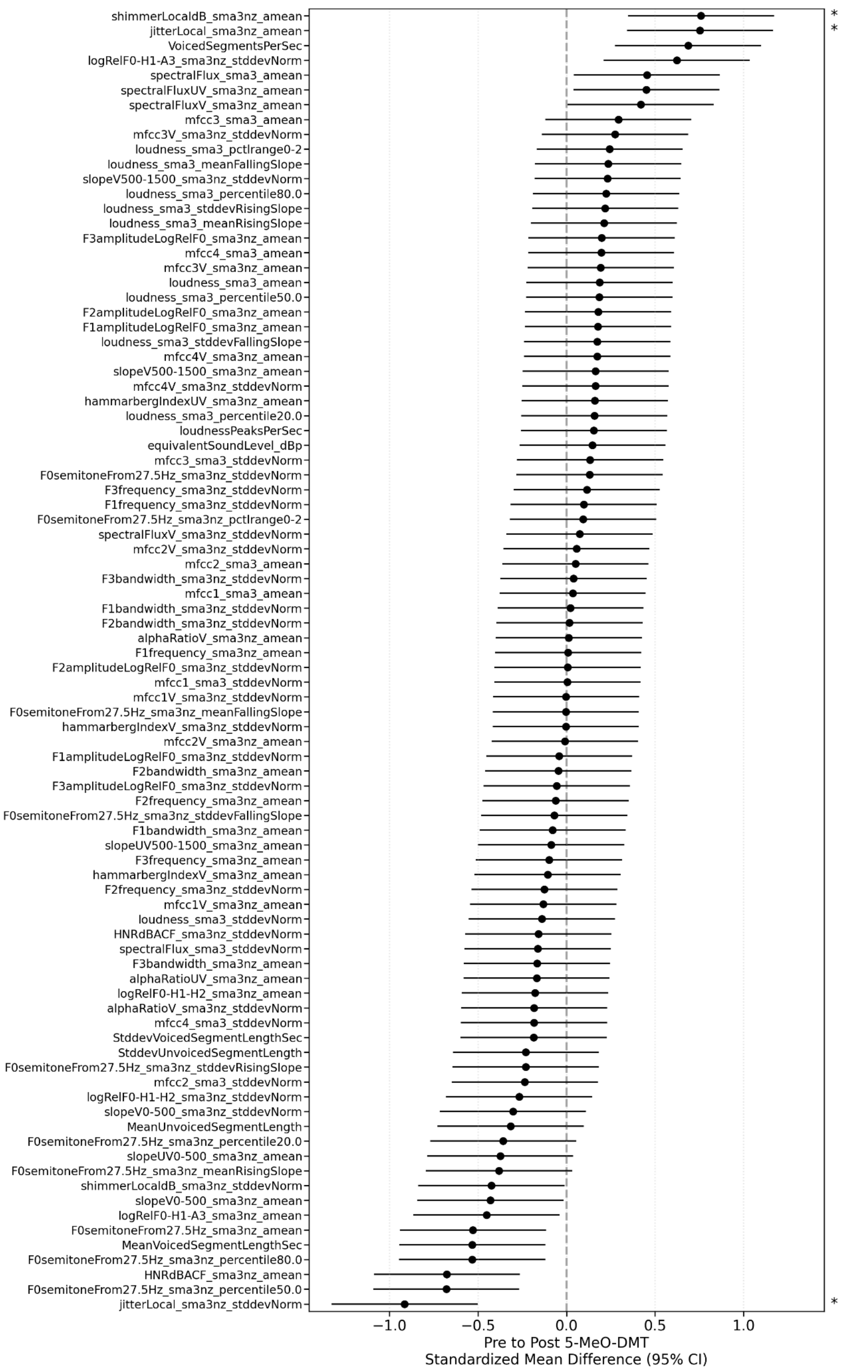


**Supplementary Figure S1**

Forest plot of standardized mean differences in 88 acoustic features extracted from voice recordings pre- and post-5-MeO-DMT administration (n = 23).

*Error bars represent 95% confidence intervals around the mean difference. Asterisks denote features with FDR-corrected p-values below 0.05 (**). Results are based on paired-sample t-tests.

| | Cognition | | | Social | | |
|---|---|---|---|---|---|---|
| **Period** | **Mean** | **SE** | **N** | **Mean** | **SE** | **N** |
| **Week -2** | -0.078 | 0.057 | 34 | -0.011 | 0.054 | 34 |
| **Week -1** | -0.091 | 0.028 | 99 | 0.064 | 0.032 | 99 |
| **Week +1** | 0.055 | 0.049 | 65 | -0.075 | 0.028 | 65 |
| **Week +2** | 0.090 | 0.048 | 31 | 0.005 | 0.040 | 31 |

**Supplementary Table S5**

*Weekly averages of cognitive and social word usage across four time periods relative to 5-MeO-DMT administration, calculated as standardized deviations from individual baselines.*

The table presents means, standard errors (SE), and sample sizes (N) for both cognitive and social LIWC categories. Means represent standard deviations from individual baselines, where positive values indicate above-typical usage and negative values indicate below-typical usage for each participant. Week -2 and Week -1 represent the two weeks before dosing, while Week +1 and Week +2 represent the two weeks after dosing. Sample sizes vary due to differences in journaling frequency across periods.

| | Cognition | | | | | Social | | | | |
|---|---|---|---|---|---|---|---|---|---|---|
| **Comparison** | **T-stat** | **P-Value** | **FDR Corr. P-Val** | **Sig.** | **Mean Diff.** | **T-stat** | **P-Value** | **FDR Corr. P-Val** | **Sig.** | **Mean Diff.** |
| **Week -2 vs Week -1** | 0.199 | 0.843 | 0.843 | ns | 0.013 | -1.187 | 0.240 | 0.358 | ns | -0.075 |
| **Week -2 vs Week +1** | -1.760 | 0.082 | 0.124 | ns | -0.133 | 1.051 | 0.298 | 0.358 | ns | 0.064 |
| **Week -2 vs Week +2** | -2.246 | 0.028 | 0.056 | ns | -0.168 | -0.235 | 0.815 | 0.815 | ns | -0.016 |
| **Week -1 vs Week +1** | -2.556 | 0.012 | 0.036 | * | -0.146 | 3.279 | 0.001 | 0.008 | ** | 0.139 |
| **Week -1 vs Week +2** | -3.229 | 0.002 | 0.013 | * | -0.181 | 1.160 | 0.250 | 0.358 | ns | 0.059 |
| **Week +1 vs Week +2** | -0.507 | 0.613 | 0.736 | ns | -0.035 | -1.653 | 0.104 | 0.311 | ns | -0.080 |

**Supplementary Table S6**

*Pairwise comparisons of cognitive and social word usage between different weeks relative to 5-MeO-DMT administration.*

For each comparison, the table presents t-statistics (T-stat), uncorrected p-values (P-Value), FDR-corrected p-values (FDR Corr. P-Val), significance levels (Sig.), and mean differences (Mean Diff.) for both cognitive and social LIWC categories. Significance levels are indicated as: $p < 0.05$ (), $p < 0.01$ (), $p < 0.001$ (), and ns for non-significant results. Mean differences represent changes in standard deviations from individual baselines between compared weeks. Notable significant changes include increased cognitive word usage from Week -1 to Weeks +1 and +2 ($p < 0.05$), and changes in social word usage between Week -1 and Week +1 ($p < 0.01$).

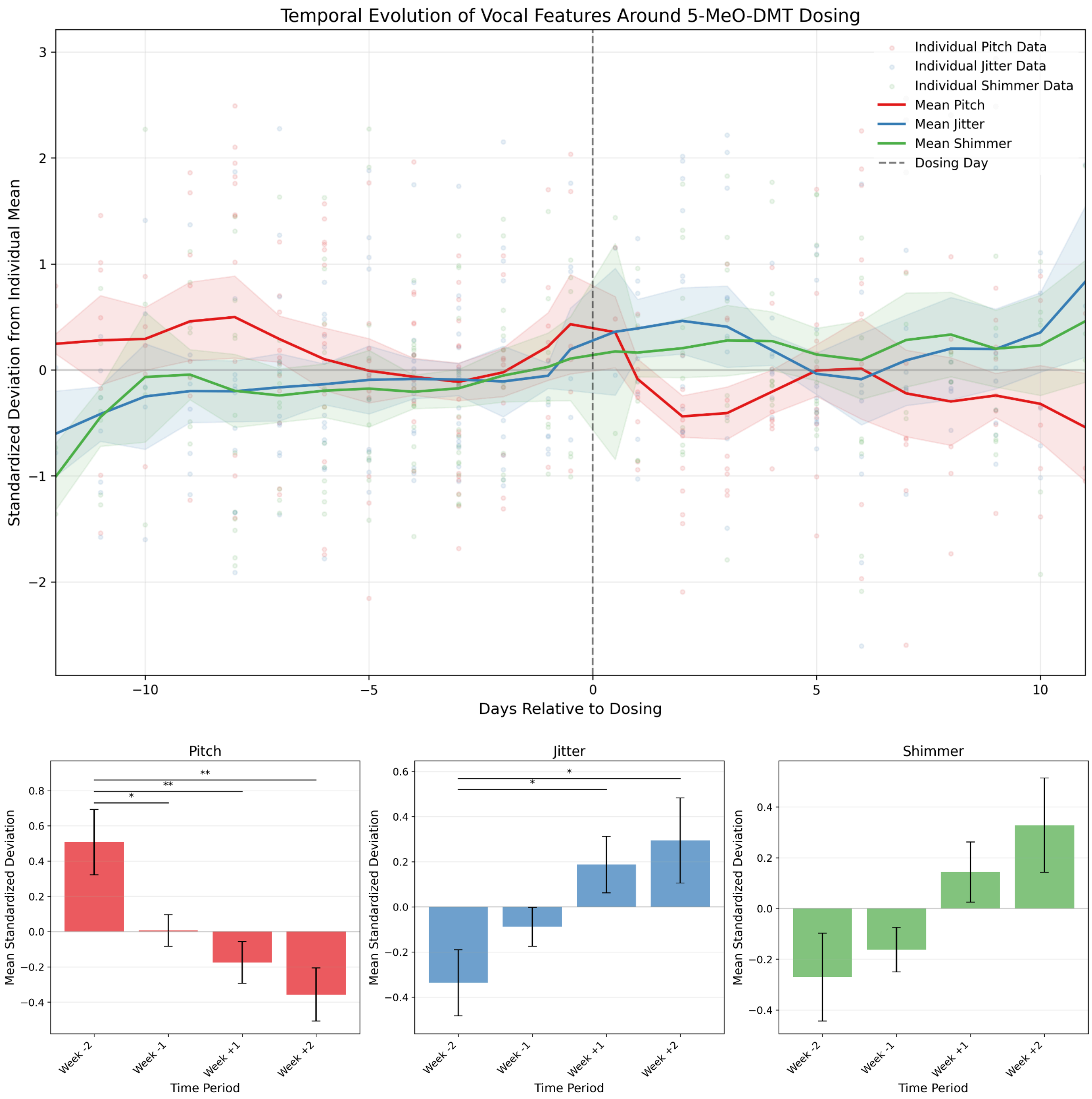


**Supplementary Figure S2**

*Temporal Evolution and Period Comparison of Standardized Vocal Features Around 5-MeO-DMT Dosing.*

(Top) Time series showing standardized deviations from individual means in pitch (red), jitter (blue), and shimmer (green) measurements from 14 days before to 14 days after dosing. Light-colored points represent individual data points, while solid lines show smoothed averages with standard error bands. Vertical dashed line indicates dosing day (Day 0). (Bottom) Bar plots depicting mean standardized deviations during each weekly period (Week -2: -14 to -7 days; Week -1: -7 to 0 days; Week +1: 0 to 7 days; Week +2: 7 to 14 days), with error bars representing standard error. Statistical significance between periods based on independent t-tests with FDR correction is indicated by * p < 0.05, p < 0.01, * p < 0.001

| Questionnaire | LIWC Feature | Coefficient | Effect_Size | Model_R2 | Model_f2 | Best_alpha |
|---|---|---|---|---|---|---|
| **ASC-DED** | sexual | 0.027 | negligible | 0.8721 | 6.818 | 46.415 |
| | quantity | 0.020 | negligible | | | |
| | focusfuture | -0.020 | negligible | | | |
| | substances | -0.018 | negligible | | | |
| | Apostro | -0.018 | negligible | | | |
| **ASC-OBN** | substances | -0.011 | negligible | 0.5083 | 1.033 | 187.381 |
| | netspeak | -0.011 | negligible | | | |
| | friend | -0.010 | negligible | | | |
| | wellness | 0.010 | negligible | | | |
| | QMark | -0.010 | negligible | | | |
| **EBI** | risk | 2.177 | large | 0.2053 | 0.258 | 1000 |
| | illness | 2.125 | large | | | |
| | netspeak | -1.906 | large | | | |
| | curiosity | -1.781 | large | | | |
| | AllPunc | -1.777 | large | | | |
| **PPS** | work | 0.331 | medium | 0.5805 | 1.383 | 284.803 |
| | Lifestyle | 0.315 | medium | | | |
| | tone_pos | 0.284 | small | | | |
| | Tone | 0.240 | small | | | |
| | swear | -0.240 | small | | | |
| **sWEMWBS** | mental | -0.098 | negligible | 0.4169 | 0.715 | 376.493 |
| | Lifestyle | 0.077 | negligible | | | |
| | Cognition | -0.069 | negligible | | | |
| | cogproc | -0.066 | negligible | | | |
| | leisure | 0.062 | negligible | | | |

**Supplementary Table S7**

*Ridge regression models predicting psychometric outcomes from pre-dosing LIWC language features.*

The table presents results from ridge regression models predicting five psychometric outcomes—psychedelic preparedness (PPS), emotional breakthrough (EBI), oceanic boundlessness (ASC-OBN), dread of ego dissolution (ASC-DED), and post-experience psychological wellbeing (sWEMWBS)—using 118 LIWC features extracted from pre-dosing journals. For each outcome, the top five predictors based on standardized coefficients are shown, along with their estimated coefficients and qualitative effect sizes (negligible, small, medium, large). Model performance metrics include cross-validated $R^2$ values (Model_$R^2$), Cohen's $f^2$ effect sizes (Model_$f^2$), and the optimal regularization parameter (Best_alpha). Where coefficient magnitudes are small despite high $R^2$ values, model interpretability may be limited.

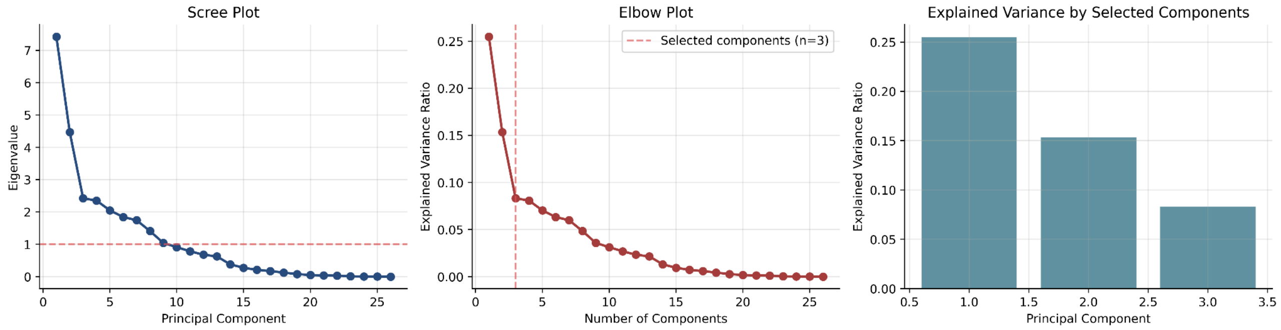


**Supplementary Figure S3**

*Variance explained by principal components extracted from pre-dosing emotion scores.*

Left: Scree plot showing eigenvalues for each principal component. Middle: Elbow plot showing cumulative explained variance ratio across components. Right: Bar plot showing individual explained variance ratio for each component.

| | PPS | | EBI | | ASC OBN | ASC DED | sWEMWBS post experience | |
|---|---|---|---|---|---|---|---|---|
| | **All PCs** | **PC1** | **All PCs** | **PC3** | **All PCs** | **All PCs** | **All PCs** | **PC1** |
| **R2** | 0.383 | 0.363 | 0.410 | 0.377 | 0.069 | 0.086 | 0.312 | 0.303 |
| **BIC** | 195.62 | 189.92 | 343.93 | 338.83 | 16.20 | 15.87 | 137.33 | 131.13 |
| **Intercept** | 124.19 | 124.19 | 257.35 | 257.35 | 0.71 | 0.28 | 29.69 | 29.69 |
| **PC1 (p-val, FDR)** | 2.326 (0.002, 0.012*) | 2.326 (0.001, 0.002**) | -8.050 (0.480, 0.817) | | 0.018 (0.398, 0.817) | -0.003 (0.875, 0.925) | 0.656 (0.005, 0.025*) | 0.656 (0.004, 0.004**) |
| **PC2 (p-val, FDR)** | -0.585 (0.490, 0.817) | | 12.199 (0.407, 0.817) | | -0.003 (0.925, 0.925) | -0.016 (0.547, 0.820) | -0.112 (0.683, 0.854) | |
| **PC3 (p-val, FDR)** | 0.524 (0.648, 0.854) | | 73.475 (0.001, 0.012*) | 73.475 (0.001, 0.002**) | 0.034 (0.359, 0.817) | 0.046 (0.209, 0.784) | 0.118 (0.752, 0.868) | |

**Supplementary Table S8**

*Multiple regression analyses examining relationships between RoBERTa-GoEmotions principal components and psychological measures.*

The table presents regression results between emotional principal components (derived from RoBERTa-GoEmotions analysis of pre-dosing journals) and five psychological measures: psychedelic preparedness (PPS), emotional breakthrough (EBI), oceanic boundlessness (ASC OBN), dread of ego dissolution (ASC DED), and post-experience psychological wellbeing (sWEMWBS). For each measure, results are shown for both full models including all principal components ("All PCs") and, where a component showed significance in the full model evaluation, additional single-component models were conducted. Model performance metrics include R-squared values indicating explained variance, Bayesian Information Criterion (BIC) for model comparison, and intercept values. For each principal component (PC1-PC3), standardized Coef.s are presented with their corresponding p-values and False Discovery Rate (FDR) corrections in parentheses.
